# Brain network dynamics during spontaneous strategy shifts and incremental task optimization

**DOI:** 10.1101/481838

**Authors:** Michele Allegra, Shima Seyed-Allaei, Nicolas W. Schuck, Daniele Amati, Alessandro Laio, Carlo Reverberi

## Abstract

With practice, humans may improve their performance in a task by either optimizing a known strategy or discovering a novel, potentially more fruitful strategy. How does the brain support these two fundamental abilities? In the present experiment, subjects performed a simple perceptual decision-making task. They could either use and progressively optimize an instructed strategy based on stimulus position, or spontaneously devise and then use a new strategy based on stimulus color. We investigated how local and long-range BOLD coherence behave during these two types of strategy learning by applying a recently developed unsupervised fMRI analysis technique that was specifically designed to probe the presence of transient correlations. Converging evidence showed that the posterior portion of the default network, i.e. the precuneus and the angular gyrus bilaterally, has a central role in the optimization of the current strategy: these regions encoded the relevant spatial information, increased the level of local coherence and the strength of connectivity with other relevant regions in the brain (e.g. visual cortex, dorsal attention network). This increase was proportional to the task optimization achieved by subjects, as measured by the reduction of reaction times, and was transiently disrupted when subjects were forced to change strategy. By contrast, the anterior portion of the default network (i.e. medial prefrontal cortex) together with rostral portion of the fronto-parietal network showed an increase in local coherence and connectivity only in subjects that would at some point spontaneously choose the new strategy. Overall, our findings shed light on the dynamic interactions between regions related with attention and with cognitive control, underlying the balance between strategy exploration and exploitation. Results suggest that the default network, far from being “shut-down” during task performance, has a pivotal role in the background exploration and monitoring of potential alternative courses of action.

## Introduction

“Practice makes perfect”, they say. By engaging long enough in any activity, major improvements in both accuracy and speed are expected. This is true for tasks as complex as playing piano and as mundane as preparing home-made tagliatelle (Ericsson & Lehmann, 1996). These improvements may happen through multiple paths. One way is incremental task optimization: while following the same solution strategy, one can optimize the implementation of the adopted (or instructed) algorithm, yielding measurable processing gains. Alternatively, one can spontaneously learn about potentially useful contingencies in the task (e.g. new stimulus-response associations), use this new information to devise a new strategy based on a different algorithm, and then switch to it, thus reaching the same task goals with greater efficiency (Badre, Kayser, & D’Esposito, 2010; Cohen, McClure, & Yu, 2007; Cole, Braver, & Meiran, 2017; Collins & Frank, 2013; Donoso, Collins, & Koechlin, 2014; Hayden, Pearson, & Platt, 2011; Heathcote, Brown, & Mewhort, 2000; Roeder & Ashby, 2016; Schuck et al., 2015; Wenke, De Houwer, De Winne, & Liefooghe, 2015).

Task optimization has been associated with a decrease of activation both in areas specialized for the task and in a set of brain regions associated to control and attentional functions (Chein & Schneider, 2005; Hampshire et al., 2016; Patel, Spreng, & Turner, 2013). The distributed nature of these effects suggests that optimisation reflects increasingly efficient routing of information across the brain. This may be understood as a modulation of neural circuits involving multiple brain regions, rather than just a local activation change. Evidence relating learning to brain-wide network changes has become available thanks to the recent development of tools for network dynamics analysis (Bassett et al., 2011; Bassett, Yang, Wymbs, & Grafton, 2015; Bassett & Mattar, 2017; Cole et al., 2013) able to capture the rich information contained in rapidly varying connectivity patterns. Nevertheless, the evidence available to date is mainly focused on motor learning, leaving open the question on how the human brain optimizes known strategies and how this relates to the discovery and implementation of new ones.

We investigated the modulation of functional networks occurring during task optimization and how this modulation interacts with spontaneous or forced strategy shifts (Schuck et al., 2015). Subjects were instructed to press one of two alternative buttons based on the spatial features of the stimuli. Even though subjects were not informed, the color of the stimuli could be used to efficiently determine the correct response. Subjects could either spontaneously discover and decide to use the new color strategy, or continue to use the instructed spatial strategy until they were explicitly told otherwise (Fig. 1).

**Figure 1.**
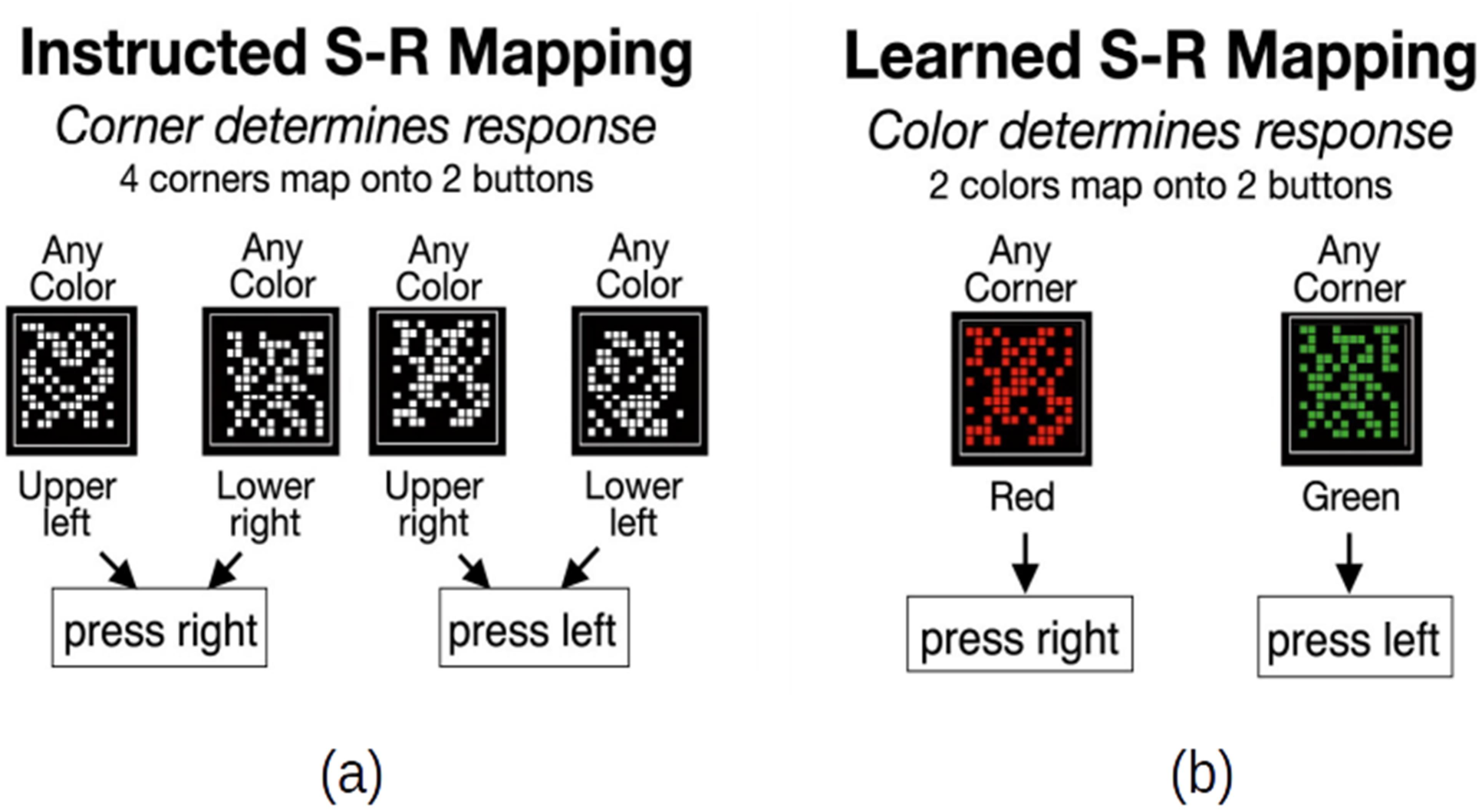
Stimulus-response mappings in the task **(a)** Instructed S-R mapping used by spatial strategy users **(b)** learned S-R mapping used by color strategy users.

The analyses were based on Coherence Density Peak Clustering (CDPC), a new fMRI analysis method developed by our group (Allegra et al., 2017). CDPC is based on Density Peak Clustering (Rodriguez & Laio, 2014), and detects the presence of clusters, sets of voxels whose BOLD time series are temporally coherent. Clusters can be reliably found even in short time windows (approximately 20 seconds). With CDPC we can identify potentially task-relevant voxels, such as those being frequently part of a cluster. Furthermore, CDPC can track the connectivity dynamics of those relevant voxels.

Overall, this new analysis method combined with our experiment allowed us to pursue multiple aims. First, we could identify, with a fully unsupervised approach, the brain regions involved in the task. Second, we traced the dynamics of local coherence and long-range connectivity associated with instructed strategy optimization. Third, we identified the BOLD signatures anticipating the discovery of a possible alternative strategy and its implementation. Fourth, we compared the changes induced by spontaneous and forced strategic shifts.

## Results

Participants performed a simple perceptual decision task (Schuck et al., 2015). They were instructed to respond manually to the position of a patch of colored dots within a square reference frame by selecting one of two responses. Participants held a button box in each hand and could press either the left or the right button. Color patches closer to the upper left or lower right corner mapped onto the left button, while patches closer to the lower left or upper right corner mapped onto the right button (Fig. 1). So there was a four-to-two stimulus-response mapping, where two opposite corners (along the diagonal) were mapped onto the same response. Participants performed 12 runs of the task, each one comprising 168 trials and lasting ~5 minutes. In runs 1 and 2, the stimulus color (red or green) was unrelated to the position of the stimulus and the response. In runs 3–12 the color had a fixed relation to the response (e.g., all upper-left and lower-right stimuli were green, the remaining ones were red, Fig. 1). Participants were not informed about this contingency but they could learn it and generate a new task strategy based on the stimulus color. Before the last two runs (11–12), participants were explicitly instructed to switch to the color strategy. For the following analyses, we considered experimental “blocks”, where 1 block = ½ run, lasting ~2.5 minutes.

### Behavioral results

Most of the behavioral results have been already reported in our previous work (Schuck et al., 2015). We briefly summarize here the findings relevant to the present work. The majority of subjects (25/36, the “spatial strategy users”) used the instructed spatial strategy over the first 20 blocks. As expected, spatial strategy users showed evidence of incremental task optimization during the first 20 blocks, as indexed by a progressive reduction of reaction times (RTs) and errors (Fig. 2a). After the instructed switch to the color strategy from block 21, RTs and errors further decreased, thus confirming the effectiveness of the color strategy. A minority of the subjects (11/36, “color users”) switched spontaneously to the color strategy before the end of the block 20. The switch point could be precisely and robustly identified by several behavioral markers (see Methods). In particular, in a fraction of the trials (ambiguous trials) the dots were centered within the square reference frame, equidistantly from all corners: in these trials, evidence of a color-based strategy comes from the number of responses that are consistent with the stimulus color (while a strategy based on stimulus position should yield essentially random responses). The fraction of color consistent-choices in color users shows an abrupt increase in the switch block (fig 2b). Before the strategy switch, color users also showed a progressive reduction in RTs and errors (Fig. 2c). This trend exhibits a transient stop just before the spontaneous switch. After the spontaneous switch to the color strategy, RTs and errors further decrease also in color users, albeit less abruptly as compared to spatial strategy users (Fig. 2d).

**Figure 2.**
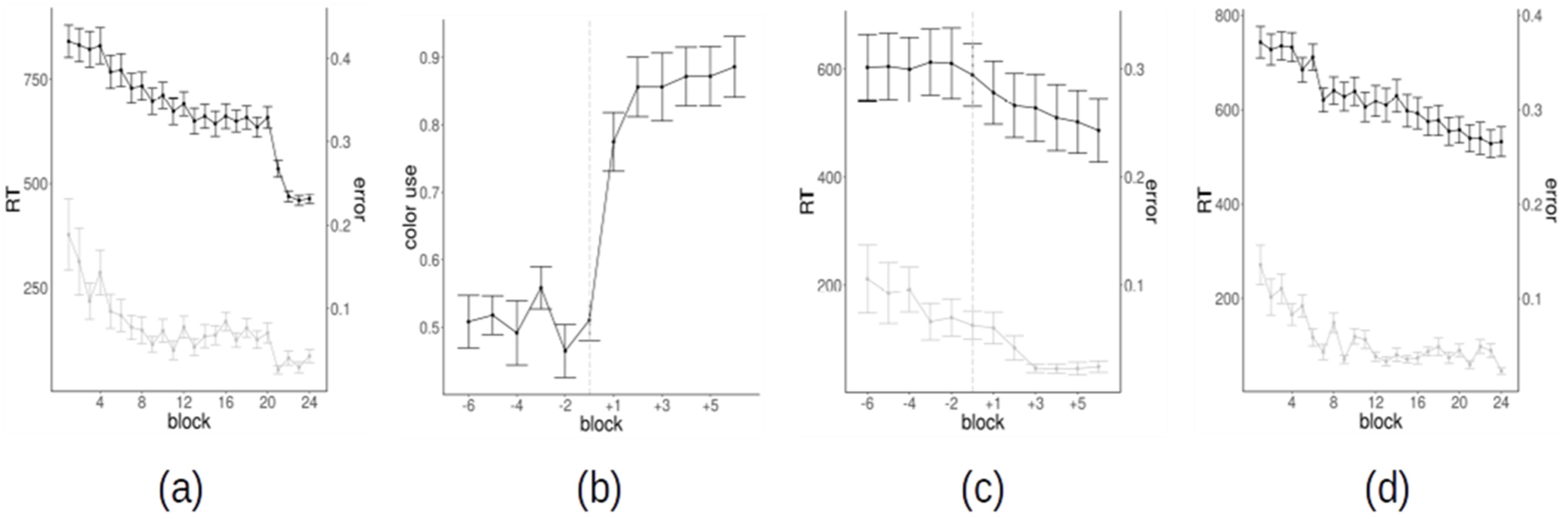
**(a)** Reaction times and error rates as a function of block for spatial strategy users **(b)** Percentage of color use (of color-consistent choices in ambiguous trials) for color strategy users as a function of block; the time series have been realigned to the switch block of each subject (−1 = switch block), which is also identified by the vertical dashed line. **(c)** Reaction times and error rates as a function of block for color strategy users; the time series are realigned to the switch block of each subject (−1 = switch block); the vertical dashed line identifies the switch **(c)** Reaction times and error rates as a function of the block for color strategy users, with time series realigned to the switch block (−1 = switch block) **(d)** Reaction times and error rates as a function of the block for color strategy users, without time series realignment.

### Overview of the neuroimaging analyses

The core of our neuroimaging analyses is based on Coherence Density Peak Clustering (CDPC) (Allegra et al., 2017).

In Fig. 3 we summarize the main steps of the analysis. In the first step, we identified all voxels potentially relevant for the task. The main idea behind CDPC is that task-related BOLD activity should organize into transient clusters of coherent activity. Given a time window, CDPC first implements a noise filter, discarding isolated voxels that are not coherent with their spatial neighbors, and then groups the remaining voxels into clusters. We applied CDPC over short (22 s) running windows in the experiment and measured how frequently any voxel is clustered, thus generating a clustering frequency (Φ) map (Fig. 3a-3c). We then found that voxels with a high value of Φ can be grouped in 22 connected regions (Fig. 3d-3f). These regions were used as a “basis” to explore the time-dependence and the subject-dependence of Φ. Finally, we focused on long-range connectivity, thus complementing the analysis on Φ, a spatially local measure. We computed a pairwise connectivity matrix between the 22 regions, measuring the frequency with which voxels in each pair of regions are clustered together (Fig. 3g-3j). By computing the variations of the connectivity matrix over blocks, we explored how behavioral changes impact the long-range connectivity.

**Figure 3.**
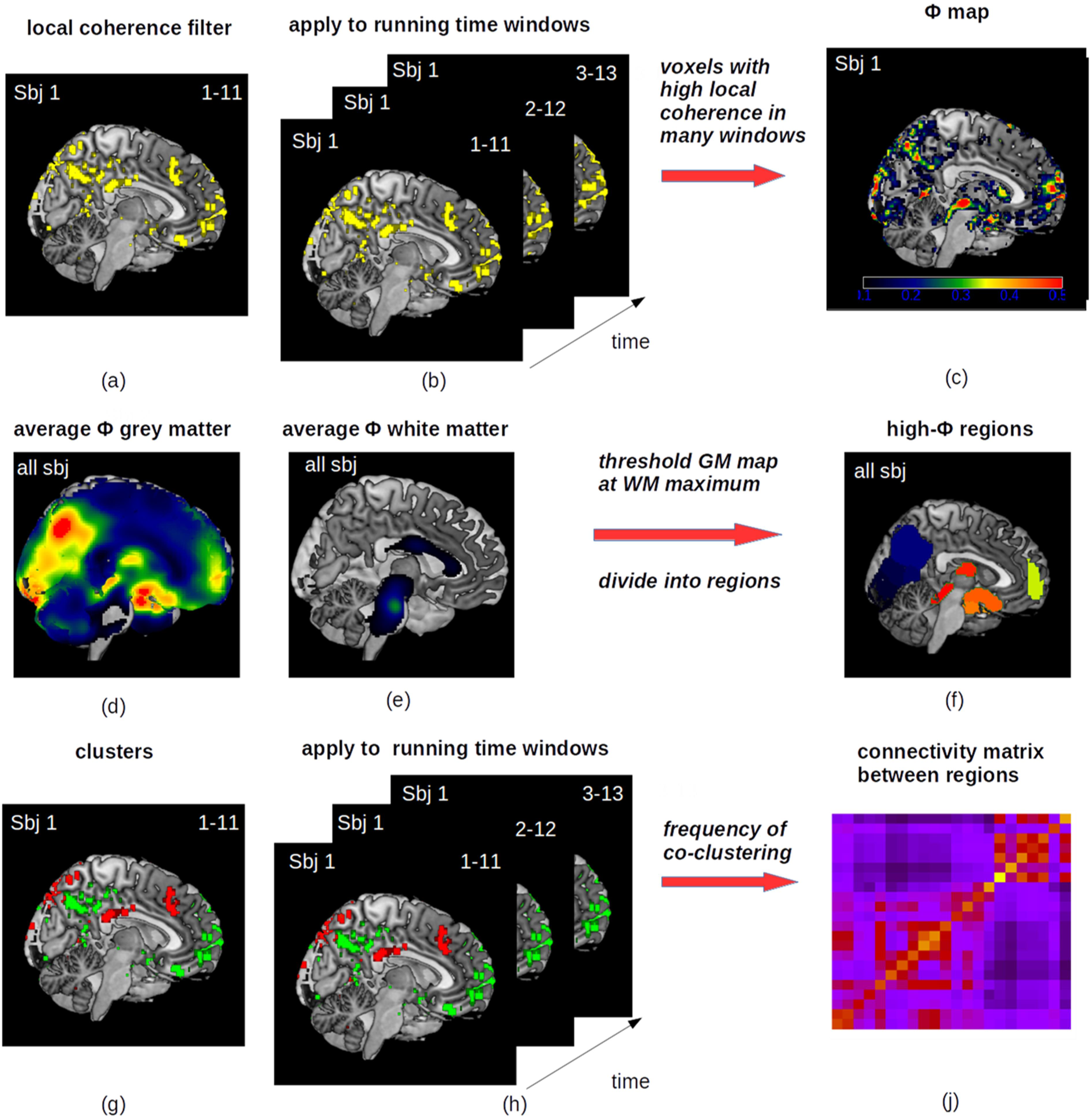
Summary of the neuroimaging analyses. In a window of 11 scans (22 s), we identify the subset of voxels that are locally coherent with at least four spatial neighbors **(a)**. Voxels surviving this local coherence filter are shown in yellow. The filtering procedure is applied separately for each subject, using sliding windows (1-11, 2-12, etc.) and including only grey matter voxels **(b)**. For each subject, we identify the frequency (fraction of time windows) with which a voxel has local coherence with its neighbors, producing a clustering frequency map Φ **(c)**. We compute an average Φ map for all subjects **(d)**. On this map, we identify voxels having significant values of average Φ. Significance is defined by computing Φ on white matter voxels **(e)**. We threshold the grey matter Φ on the maximum value found in white matter. The above-threshold voxels are divided in 22 regions around each peak of the average Φ. Different regions are shown in different colors **(f)**. Finally, we compute a connectivity matrix between high-Φ regions. For each subject separately, in each time window of 11 scans we consider the voxels surviving the spatial filter and we divide them into different coherent clusters based on density peak clustering **(g)**. Voxels assigned to two different clusters are shown in red and green respectively. We compute the clusters in all time windows **(h)** and define a pairwise connectivity between two regions by computing with which frequency voxels belonging to two regions assigned to the same cluster **(j)**.

### Identification of the relevant regions

In the following analyses, we considered fMRI data from 35 participants. One subject was excluded because the field of view did not cover the whole brain. For each subject, and for each of the 24 independent blocks, we applied CDPC to grey matter voxels as identified by tissue segmentation. CPDC was applied on sliding windows of 11 volumes (22 seconds), the same timescale for which the method was validated. For each subject and each block, we computed a clustering frequency map Φ_*i*_ that measures the fraction of time windows within a block in which a voxel is coherent with its spatial neighbors and hence clustered.

The Φ_*i*_ maps are consistent across subjects: averaging Φ_*i*_ maps over all blocks and performing spatial smoothing (9 mm FWHM), we obtained a between-subject correlation of .623 (SD = .003). A comparison between the average Φ_*i*_ maps for color and spatial strategy users failed to detect any significant difference. We thus averaged Φ_*i*_ maps over all subjects, and we obtained a 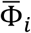 map reporting the average clustering frequency across subjects and blocks. Areas with high 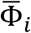 represent voxels where a coherent signal is consistently found over different blocks and subjects. To estimate which values 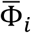 could be expected in the absence of a real task-related signal, the same procedure was carried on for voxels in the white matter. We assumed that task processing did not modify Φ in the white matter. To identify potentially task-relevant voxels we conservatively thresholded the 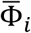 map of the grey matter with the maximum value 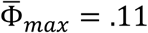 observed in the white matter. In Fig. 4 we show the resulting thresholded 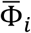 map for gray matter. We grouped spatially contiguous voxels above the white matter threshold in regions around each peak in the 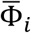 map, thus obtaining 22 regions (Tab. 1). These high-Φ regions are distributed throughout the brain, including areas in the occipital cortex, parietal cortex, prefrontal cortex, temporal cortex, thalamus, and mesencephalon. These 22 regions will be used in the following as a “basis”, common to all subjects and all runs, for describing the time- and subject-dependence of the results.

**Figure 4.**
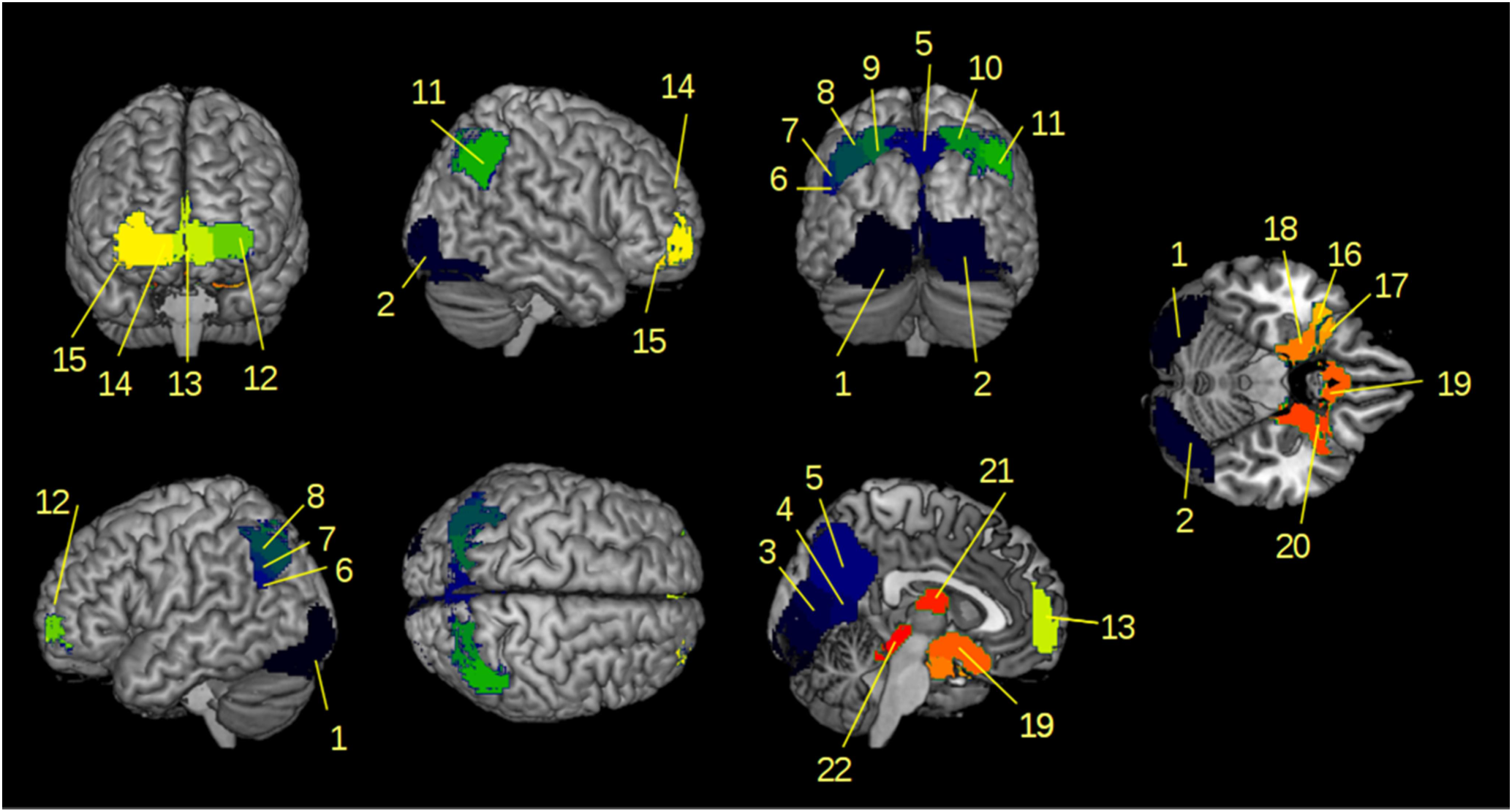
Voxels in gray matter showing an average clustering frequency (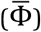) higher than the maximum clustering frequency in white matter (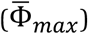). Voxels have been grouped in 22 regions around each peak of the 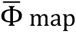 map.

**Table 1:** Summary information about the 22 high-Φ regions, including brain location (main AAL region and Brodmann area, MNI coordinates of the region center), size (number of voxels), and the short name used in figures.

| # | Brain Region (AAL) | Brodmann Area | Size | Coordinates |  |  | Short name |
| --- | --- | --- | --- | --- | --- | --- | --- |
|  |  |  |  | x | y | z |  |
| 1 | Left Lingual Gyrus | 18 | 2212 | -24 | -86 | -13 | 1 Occipital |
| 2 | Right Lingual Gyrus | 18 | 2799 | 21 | -84 | -9 | 2 Occipital |
| 3 | Calcarine Sulcus | 17 | 528 | 1 | -77 | 11 | 3 Occipital |
| 4 | Calcarine Sulcus | 17 | 310 | 0 | -61 | 9 | 4 Occipital |
| 5 | Left Precuneus | 7 | 2407 | 0 | -65 | 38 | 5 Parietal |
| 6 | Left Angular Gyrus | 39 | 90 | -50 | -60 | 27 | 6 Parietal |
| 7 | Left Angular Gyrus | 39 | 144 | -47 | -62 | 33 | 7 Parietal |
| 8 | Left Angular Gyrus | 39 | 586 | -39 | -65 | 43 | 8 Parietal |
| 9 | Left Superior Parietal Lobule | 7 | 455 | -24 | -67 | 50 | 9 Parietal |
| 10 | Right Superior Parietal Lobule | 7 | 748 | 26 | -68 | 48 | 10 Parietal |
| 11 | Right Angular Gyrus | 39 | 757 | 45 | -57 | 44 | 11 Parietal |
| 12 | Left Superior Frontal Gyrus, orbital part | 10 | 440 | -25 | 58 | -3 | 12 Frontal |
| 13 | Left Medial Orbitofrontal Cortex | 10 | 815 | -2 | 58 | 0 | 13 Frontal |
| 14 | Right Superior Frontal Gyrus, orbital part | 11 | 748 | 26 | 58 | -5 | 14 Frontal |
| 15 | Right Middle Frontal Gyrus, orbital part | 11 | 80 | 36 | 46 | -14 | 15 Frontal |
| 16 | Left Superior Temporal Pole | 38 | 150 | -34 | 7 | -23 | 16 Temporal |
| 17 | Left Superior Temporal Pole | 38 | 79 | -27 | 11 | -21 | 17 Temporal |
| 18 | Left Hippocampus | 28 | 475 | -17 | -3 | -20 | 18 Temporal |
| 19 | Caudate |  | 1039 | 0 | 8 | -15 | 19 Caudate |
| 20 | Right Hippocampus | 34 | 779 | 21 | 0 | -20 | 20 Temporal |
| 21 | Right Thalamus |  | 330 | -1 | -8 | 14 | 21 Thalamus |
| 22 | Mesencephalon |  | 222 | 1 | -32 | -9 | 22 Mesencephalon |

Since CDPC is an unsupervised technique, the high level of local coherent activity in the identified high-Φ regions is not necessarily related to the task. To verify which of the high-Φ regions are indeed task-related, we used multiple approaches. First, we compared CDPC results with the supervised, multivariate pattern analysis (MVPA) performed on the same regions (see Methods, and for details see also Schuck et al., 2015). For each subject and each block, MVPA produced accuracy maps assessing whether local activity patterns represented relevant features of the stimuli, namely to which corner the patch is nearest. For each high-Φ region, we tested whether the average accuracy of the multivariate classifier was above the chance level. We found that several regions in the occipital, parietal and prefrontal cortex encoded spatial information (*p* < .05, FDR corrected, Fig. 5). A second approach to investigate task-relevance is to test whether the identified regions had a different average activation during task performance as compared to a resting baseline. Almost all regions showed either significant activation or significant de-activation (*p* < .05, FDR corrected, Fig. 5). Regions more active than baseline are medial and lateral occipital cortex bilaterally (1–2), and the superior parietal lobule bilaterally (9–10). All remaining areas, including occipital (3–4), central and lateral parietal (5–8, 11), prefrontal (12–15), temporal (16–20) are deactivated. Taken together, these findings suggest that most of the identified regions are likely related to the task. Further converging evidence is provided by the subsequent analyses.

**Figure 5.**
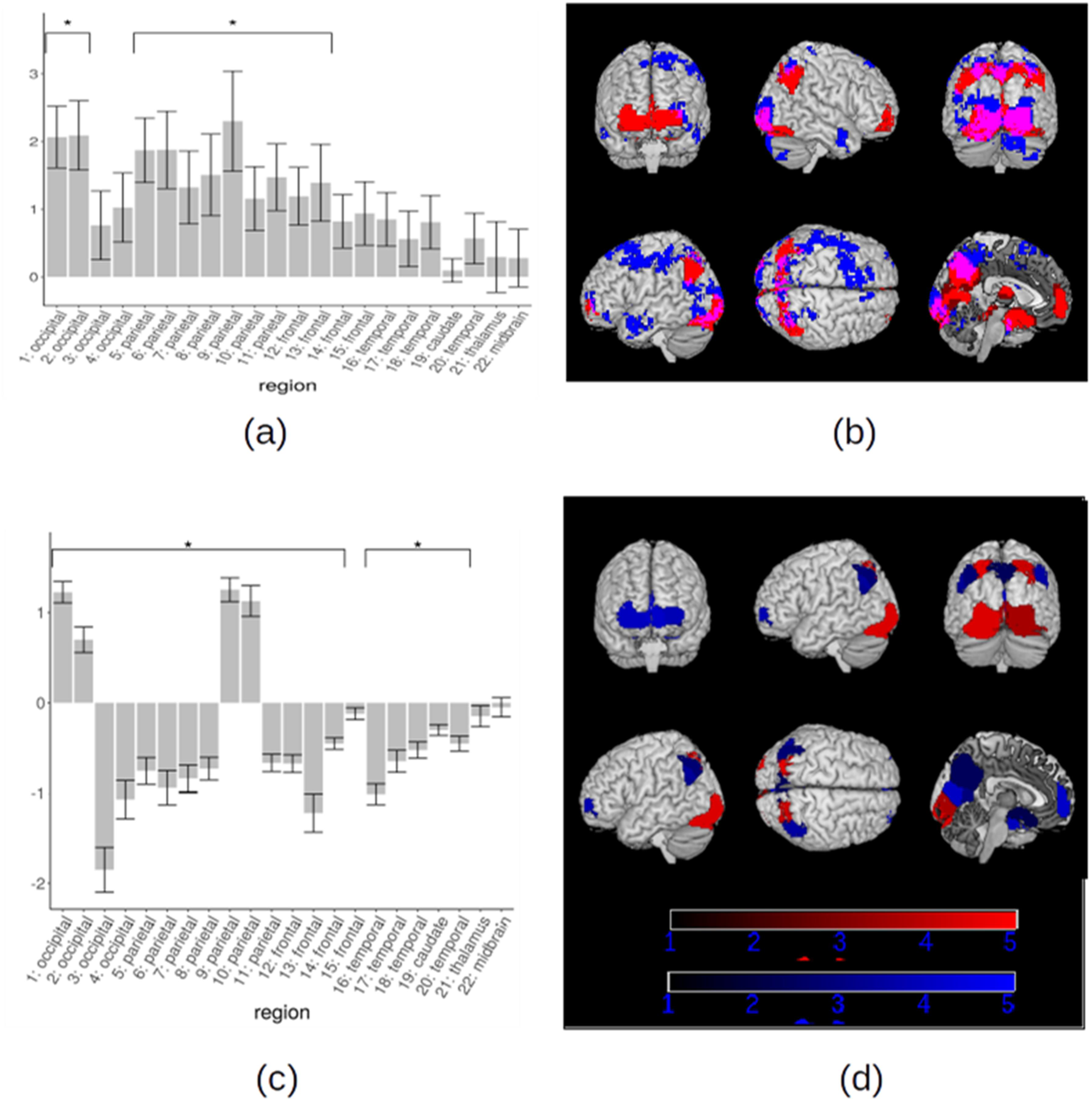
**(a)** MVPA analysis on the spatial position of the stimulus. Mean accuracy of the multivariate classifier in CDPC regions for all users. We mark with an asterisk (*) regions with significant mean corner accuracy (FDR correction at α = .05, *T*-test). **(b)** Overlap (purple) between CDPC regions (red) and voxels with significant spatial representation (cluster-wise FWE correction at α = .05, *T*-test) for spatial strategy users (blue). **(c)** Mean activation for task versus rest contrast for all subjects. We mark with an asterisk (*) regions with significant activation or deactivation (FDR correction at α = .05, *T*-test). **(d)** Regions with significant activation (red) or deactivation (blue) for task versus rest contrast as revealed by GLM analysis for all subjects (color scale represents significance, −log(*p*), 5 corresponds to *p* = 10^-5^)

### Temporal dynamics of the clustering frequency (Φ)

Having identified the set of high-Φ regions, we focused on differences in Φ related to behavioral changes across time and across subjects. Across time, we may observe a qualitative change of the brain regions showing a high level of coherence. Alternatively, high-Φ regions may remain the same but the level of Φ in the regions may systematically change.

We found that the Φ maps obtained were similar for different blocks within single subjects. We computed a block similarity for each subject by computing the Pearson correlation between all pairs of block Φ maps within each subject and averaging over blocks. Over all subjects, we obtained an average similarity of .64 (SD = .09), indicating an overall qualitative stability of the brain regions involved during the experiment. Consistently with this finding, the set of regions with significant coherence appeared stable during the experiment. For both color and spatial strategy users, we considered the voxels above the threshold 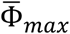 in each block. Results are reported in Fig. 6. High-Φ regions do not qualitatively change during the experiment. No regions disappeared, and no new regions appeared.

**Figure 6.**
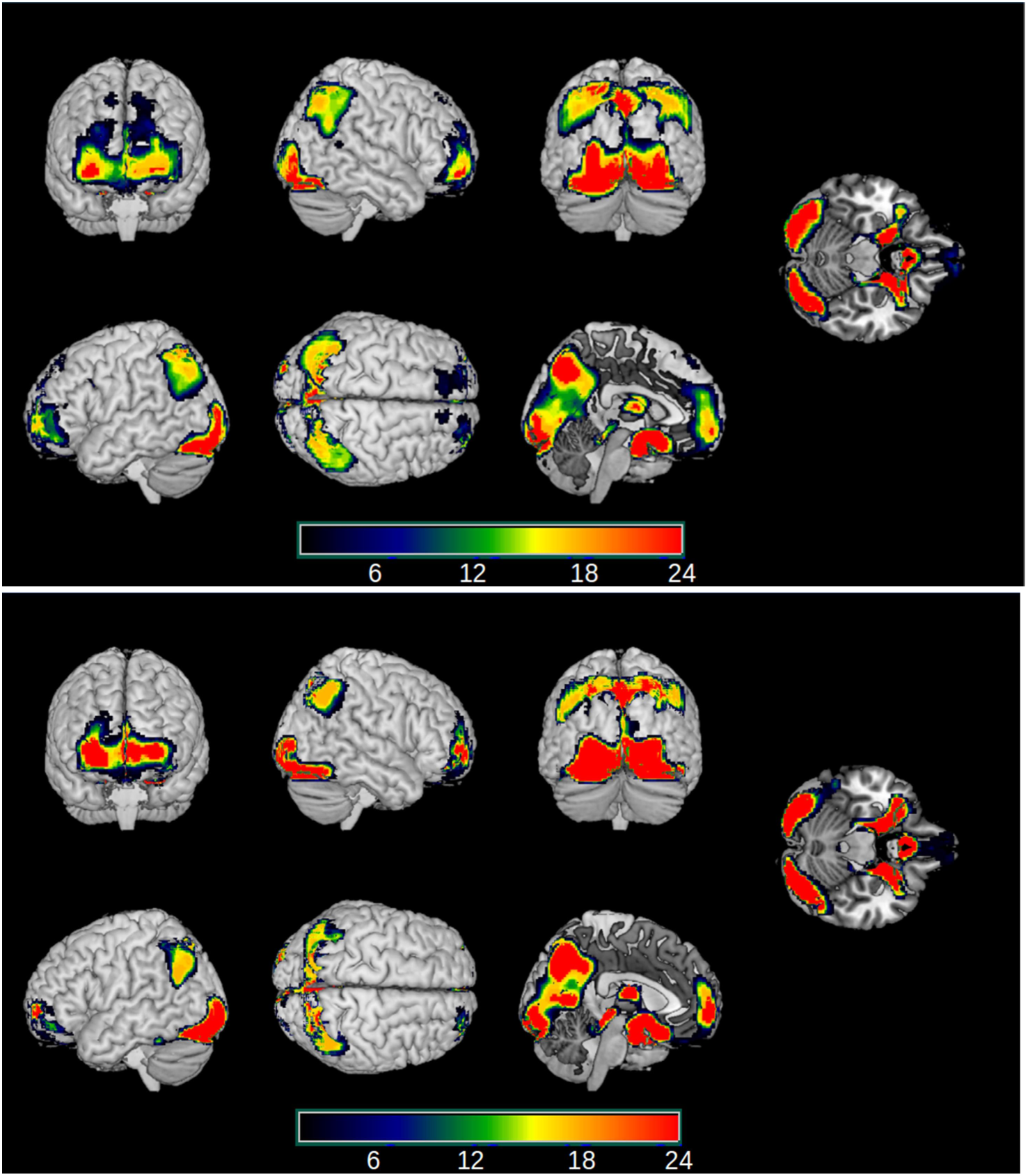
Stability of the high-Φ regions during the experiment. For each block separately, we considered the Φ for that block, averaged over subjects (top: corner users; bottom: color users), and identified the voxels with average Φ higher than the maximum value 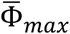 found in the white matter. Most voxels pass the threshold for almost all blocks. Here we show the conjunction map, representing the number of blocks for which a voxel passes the threshold.

The lack of qualitative change, however, does not imply that Φ remains constant over time. In fact, quantitative variations of Φ were observed. We first analyzed the subjects who used the spatial strategy up to block 20 and switched to color strategy in blocks 21–24, after receiving explicit instructions. We computed the average Φ in each region as a function of the block. The variations of Φ across blocks were different between regions. In regions 3,4 (occipital) and 5,6,7,9,11 (lateral and medial parietal) we observed three phenomena, as shown in Fig. 7 (a-e). First, in blocks from 1 to 20, i.e. when subjects used and improved the spatial strategy, Φ rose. Second, after the switch to the color strategy, Φ suddenly dropped. Third, Φ underwent a fast recovery, so that the measure was back to the pre-switch level after just one block (~2.5 min). Regions 9,10 (lateral superior parietal) showed both the increase and the drop but did not show the recovery. Regions 15–20 (temporal lobe) showed the increase, but no drop. Finally, the remaining regions, including 1,2 (occipital lobe), 12–15 (prefrontal cortex) and 21,22 (thalamus and mesencephalon) show little variation of Φ across time. The effects corresponding to the increase, drop and recovery are summarized for all regions in Fig. 8 and in Tab. 2.

**Figure 7.**
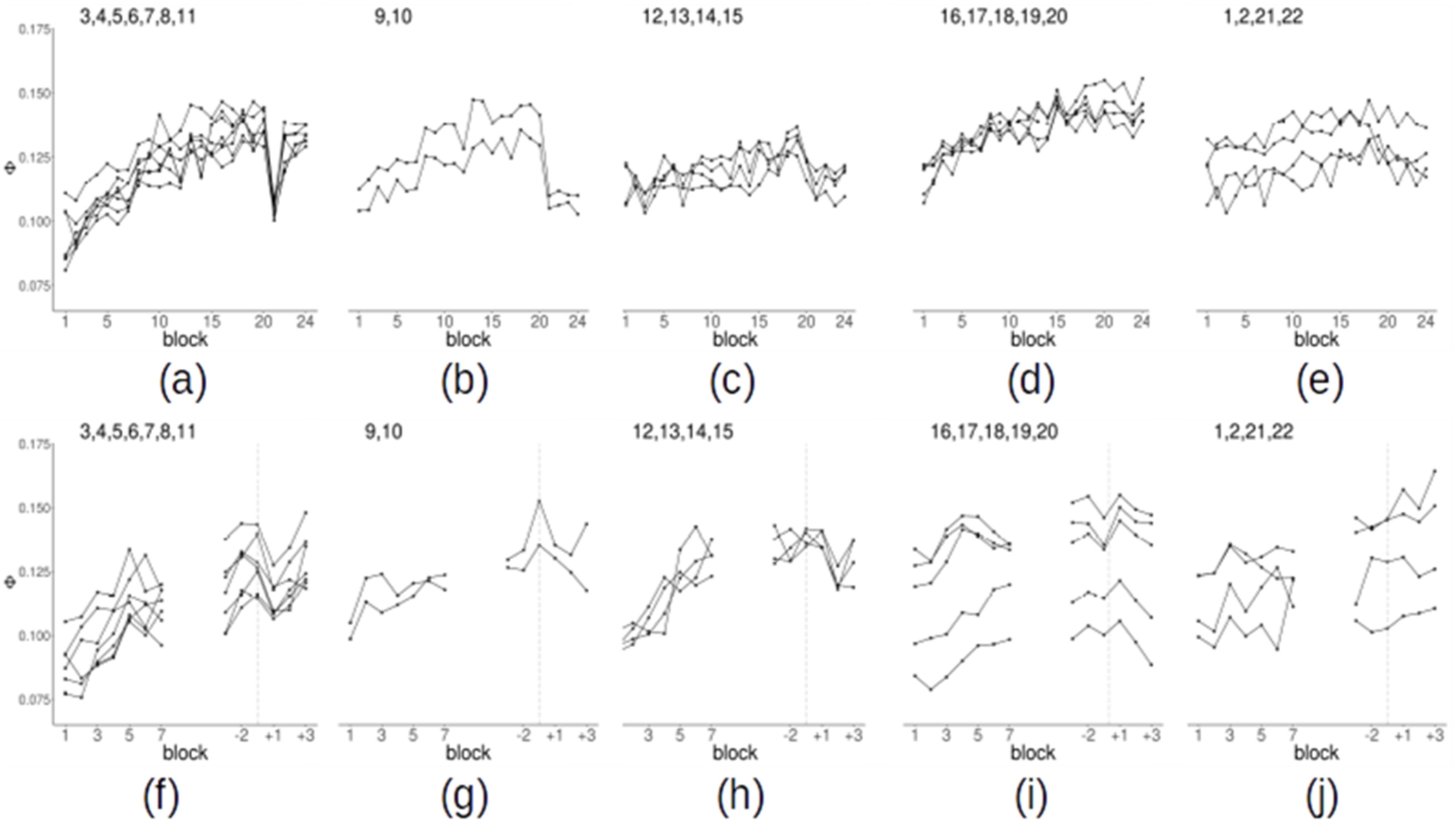
**(a-e)** Change of the average clustering frequency (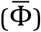) as a function of the block for spatial strategy users. **(a)** Occipital regions 3–4 and parietal regions 5–9,11 **(b)** parietal regions 9–10 (c) frontal regions 12–15 **(d)** temporal regions 16–20 **(e)** occipital regions 1–2, thalamus 21, midbrain 22. Points are mean over subjects. **(f-j)** Change of the clustering frequency as a function of the block for color strategy users. On the left of each plot, we show the first 7 blocks. On the right, we show the 6 blocks around the switch after realigning the time series of each subject to the individual switch blocks (−1 is the block in which the subjects spontaneously switch strategy, +1 the subsequent block). The vertical dashed line identifies the switch.

**Figure 8.**
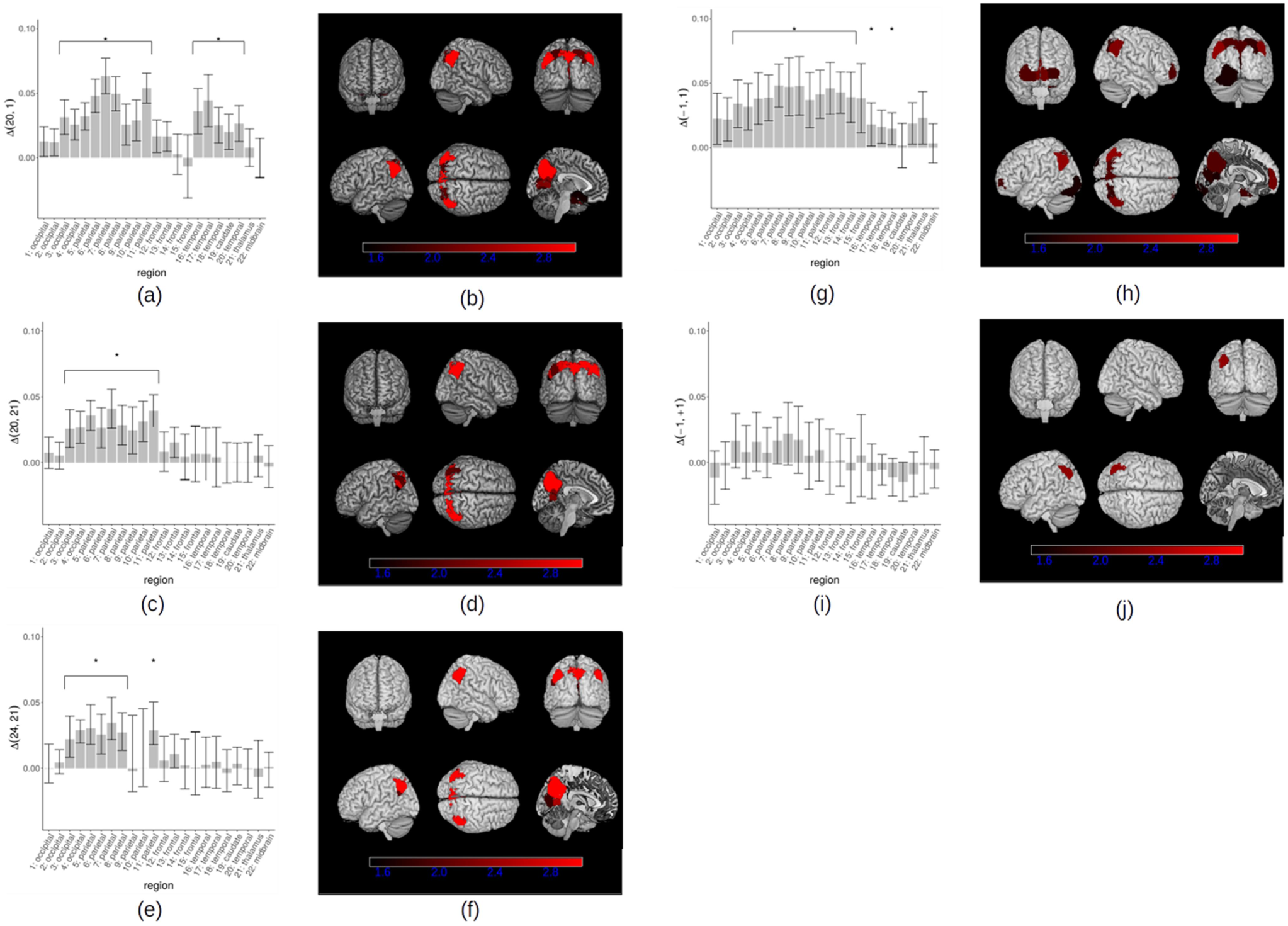
Clustering frequency (Φ) variations in high-Φ regions. **(a-b)** Difference of clustering frequency between block 20 and block 1 in spatial strategy users. **(c-d)** Difference of clustering frequency in the high-Φ regions between block 20 and block 21 in spatial strategy users. **(e-f)** Difference of frequency variation in the high-Φ regions between block 24 and block 21 in spatial strategy users. **(g-h)** Difference of clustering frequency between the block in which subjects spontaneously switched strategy (block “-1”) and the first block (block “1”) in color strategy users. **(i-j)** Difference of clustering frequency in the high-Φ regions between the block in which the subject spontaneously switches strategy (block “-1”) and the following block (block “+1”) for color strategy users. Column bars represent the average over subjects, error bars the standard error. We mark with an asterisk (*) regions where Φ increases significantly (Wilcoxon test, FDR correction at α = .05). In the rendering, we show regions with *p* < 0.05 (unc.), the colorbar represents −log(*p*)

As mentioned, several regions showed at the same time an increase in Φ during task optimization and a decrease upon strategy change (Fig. 8, Tab. 2). In these regions, the dynamics of Φ in blocks 1–20 and 20–21 is what would be expected for a variable correlated to the optimization of the spatial strategy: a gradual increase during learning of a specific strategy and a drop when such strategy is abandoned. In order to further explore this possibility, we evaluated the correlation between Φ and participants’ reaction times (RTs). Overall, the average RTs in blocks 1–20 are negatively correlated with Φ averaged over all regions (Fig. 9). We computed the correlation between Φ and RTs in each region and then averaged over subjects. The statistical significance of the resulting average correlation was assessed by using a permutation approach, re-computing the Pearson coefficient for 10,000 random permutations of the blocks. We found that the increase in Φ in most regions was significantly correlated with the decrease of RTs (*p* < .05, FDR corrected for multiple comparisons). The strongest effects were in the parietal, occipital and temporal regions (Fig. 9). If the observed negative correlation were just due to an unspecific, time-dependent, increase of Φ we would expect to find the same relation also in brain areas outside the 22 high-Φ regions examined. We assessed whether the negative correlation is present also in other brain regions by using a standard whole-brain atlas parcellating the brain in 268 regions (Finn et al., 2015). Only 28 regions out of 268 showed a significant negative correlation (*p* < .05, uncorrected). These regions are located in the parietal, precuneus and prefrontal cortex, with a large overlap with high-Φ regions. Finally, we checked whether the increase of Φ would be present even in an experiment in which task optimization is not expected to occur. If the increase of Φ were just due to artifacts or physiological noise, the increase should be observed irrespectively of whether some learning occurs in the experiment or not. We repeated the same analysis procedure on an experiment (Reverberi et al., 2018) where 15 subjects performed 7 runs of a simple language task (naming of common objects) not expected to trigger any learning. We evaluated the dynamics of Φ as a function of the run in the same regions showing a time-dependent increase in the present study. We did not observe any evidence for an increasing Φ in the language experiment (region 1: *p* = .01 uncorrected; region 6: *p* = .04 uncorrected; all other *p*s > .1, uncorrected). Thus, the increase of Φ in the present study is likely related to the task.

We carried out on color users an analysis similar to the one performed on spatial strategy users. It should be noticed that in color users the timing of the strategy change was not fixed as in spatial strategy users, but it was variable from subject to subject. Thus, we considered the increase of Φ from block 1 to the block of the strategy change. We again observed a significant increase of Φ in the regions already reported for spatial strategy users (Fig. 7 f-j). In contrast with spatial strategy users, however, in color users such an increase was observed also in prefrontal cortex (regions 12–15). Furthermore, color users showed no sudden decrease in Φ between blocks 20 and 21. This was expected given that color-users did not switch strategy at that point in time. Relatedly, we explored the presence of a decrease of Φ between the block in which subjects spontaneously changed strategy and the following one (equivalent to blocks 20/21 in spatial strategy users). We observed a decrease of Φ in the same regions that showed an effect in the spatial strategy users. The effect, however, is considerably weaker compared to the spatial strategy users. In fact, results are not significant after FDR correction; the largest effect is observed for region 8 (left parietal) with *p* = .005 (uncorrected). The weakness of the effect may be related to the reduced sample of color users (11 instead of 24 subjects). More importantly, the transition between the two strategies, when non-instructed, is likely to be gradual and, for a short period of time, the two strategies might be used simultaneously.

**Figure 9.**
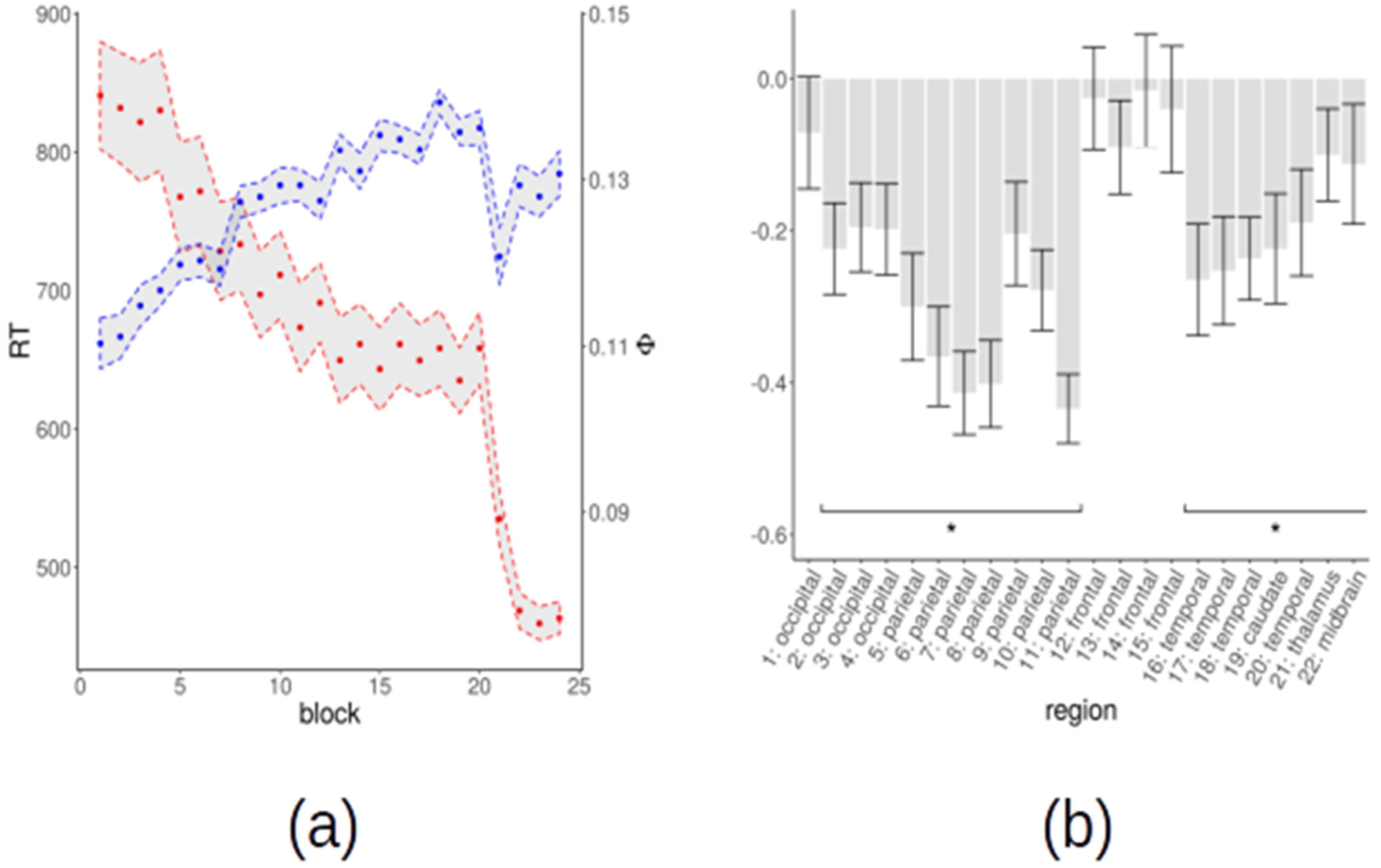
**(a)** Average reaction times (red) and average clustering frequency (blue) in high-Φ regions as a function of the block for spatial strategy users. Points are the mean over subjects, shaded regions ± the standard error. For each subject, reaction times are averaged over all trials in a block, and Φ is averaged over all regions. **(b)** Pearson correlation between Φ and reaction times in high-Φ regions for spatial strategy users. We mark with an asterisk the regions that have a significant correlation (permutation test, *p* < .05).

### Cluster-based connectivity analysis

Clustering frequency (Φ) maps reveal how often a voxel is involved in a coherent activity in a time window of approximately 20s and in a spatially local neighborhood. A group of voxels generally shows high coherence not only with voxels in the spatial neighborhood but also at large distance. Φ maps, however, do not directly measure long-distance coherence, neither can reveal which regions, among those with high-Φ, have mutually coherent time series, i.e. are connected. To quantify long-range coherence effects we measured the frequency with which voxel pairs in two regions are assigned to the same cluster, weighted by a measure of the robustness of cluster assignment (Methods). By computing these values for all regions we built a clustering-based connectivity matrix (Fig. 10). The diagonal terms of the matrix represent a measure of the within-region coherence. By contrast, the off-diagonal terms represent the coherence or connectivity between regions.

**Figure 10.**
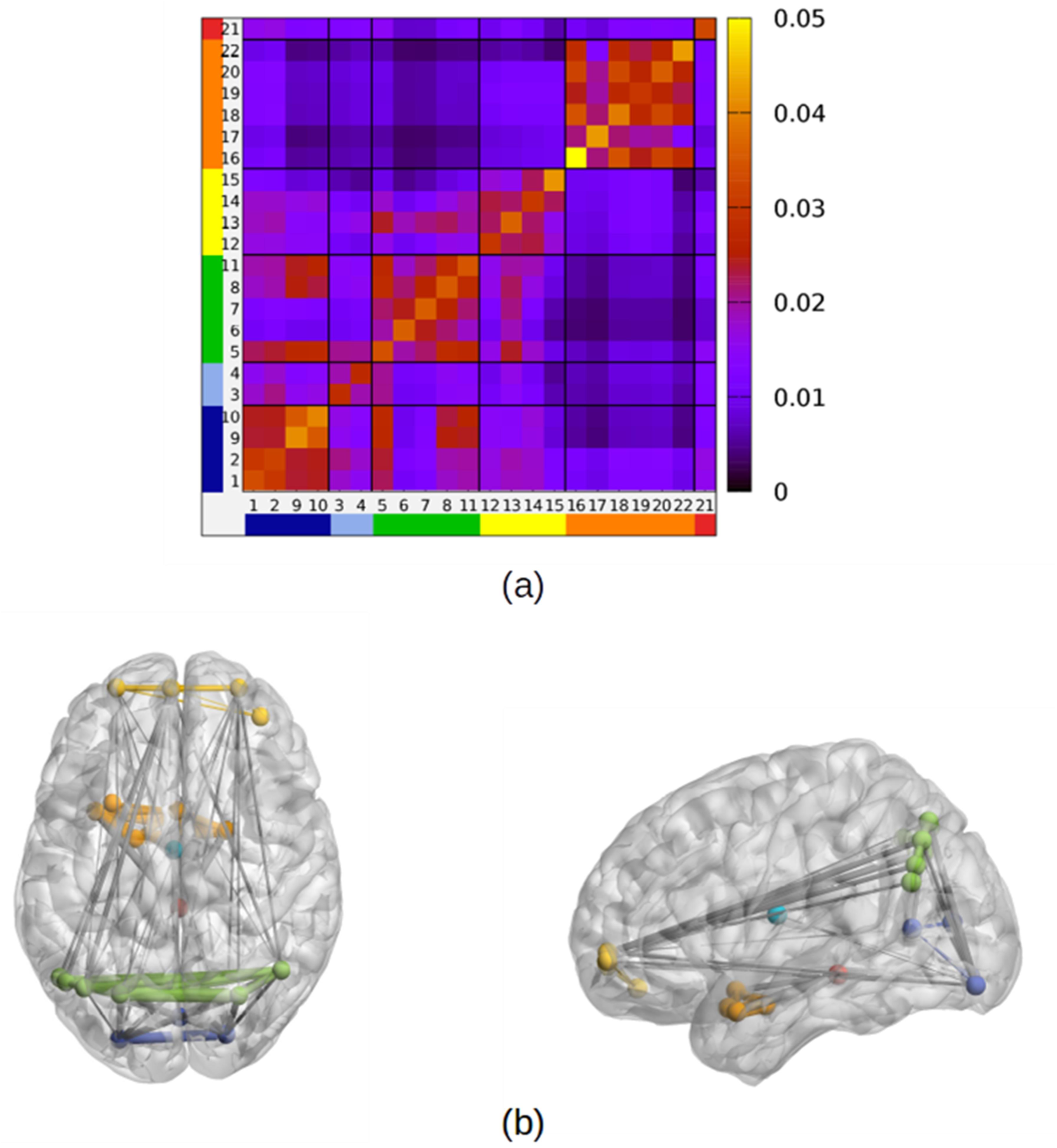
**(a)** Connectivity matrix between high-Φ regions, averaged over all subjects. The matrix element value corresponds to the value of *N*_*ab*_. Regions have been assigned to 6 modules according to a modularity maximization algorithm. The 6 modules are separated by black lines and are shown in different colors: occipito-parietal module (OP, regions 1–2,9–10), occipital module (O, regions 3–4), parietal module (P, regions 5–8,11), frontal module (F, regions 12–15), temporal module (T, regions 16–20,22), thalamus (Th, region 21). **(b)** the 50% strongest links in the average network in axial and sagittal view. Nodes assigned to different modules are shown in different colors (blue) O module (cyan) OP module (green) p module (yellow) F module (orange) T module (red) Th module.

Like the Φ-maps, the connectivity matrices obtained are similar for different blocks within a single subject (*r* = .79, SD = .06) and, upon averaging over blocks, across different subjects (*r* = .65, SD = .15). In general, the within-region coherence (average *N_aa_* = .035, SD= .006) is higher than the between-region coherence (average *N_ab_* = .008, SD= .010). Nevertheless, we found strong long-range links between subsets of regions. In Fig. 10 we show the connectivity matrix averaged over blocks and subjects, and the 50% strongest links. We used the popular Louvain modularity optimization method (Blondel, Guillaume, Lambiotte, & Lefebvre, 2008) to assign regions to subnetworks or modules. Regions within a module display higher mutual connectivity. The optimal partition identifies 6 modules. The modules are frontal (F), parietal (P), occipito-parietal (OP), occipital (O), temporal (T) and thalamus (Fig. 10). Furthermore, the precuneus (region 5) acts as a hub connecting the four fronto-parieto-occipital modules, thus showing high connectivity with all of them. More in detail, all regions in the anterior frontal cortex (12–15) are assigned to the frontal module. The regions in the temporal cortex (16–20) form the temporal module including also the midbrain (22). The thalamus (21) is not connected with other regions and forms a module alone. The occipital and parietal regions are split into 3 modules. The parietal module includes the precuneus (5) and the inferior lateral parietal regions (6–8,11). The occipito-parietal module joins two occipital regions (1,2) and the superior lateral parietal regions (9,10). Finally, two occipital regions (3,4) form the occipital module.

By using an approach similar to the one used for Φ, we explored variations of the connectivity network in time. We first analyzed spatial strategy users (Fig. 11). We observed an increase of connectivity centered on the medial and lateral parietal lobe involving namely the parieto-parietal, parieto-occipital and parieto-frontal links (*p* < 0.05 uncorrected, Wilcoxon test; links with *p* < 0.01 are also FDR corrected). Upon switch to color strategy (block 21), the strength of the links mainly centered on the parietal lobe decreased and then, as for clustering frequency, the same links showed a rebound to the connectivity level reached before the switch (*p* < 0.05 uncorrected, Wilcoxon test). In Fig. 11a-c, we show the links with an increase between block 1 and block 20, those with a decrease between block 20 and 21, and those with an increase between block 21 and 24 (*p* < .05 uncorrected, Wilcoxon test). It is apparent that there is an increase between block 1 and block 20 of the connectivity within the p module, and between the p module and the OP, O, and F modules. To improve statistical sensitivity, we performed the same tests focusing on module-wise connectivity. We averaged over all pairs of links between regions assigned to two modules: for example by averaging all links between module P regions and module OP regions we obtained P-OP connectivity (Fig. 12a). The P-P, P-OP, P-O, P-F, and O-F links have a significant increase in between block 1 and block 20 (*p* < .05 FDR corrected, Wilcoxon test) and a significant decrease between block 20 and block 21 (*p* < .05 FDR corrected, Wilcoxon test).

**Figure 11.**
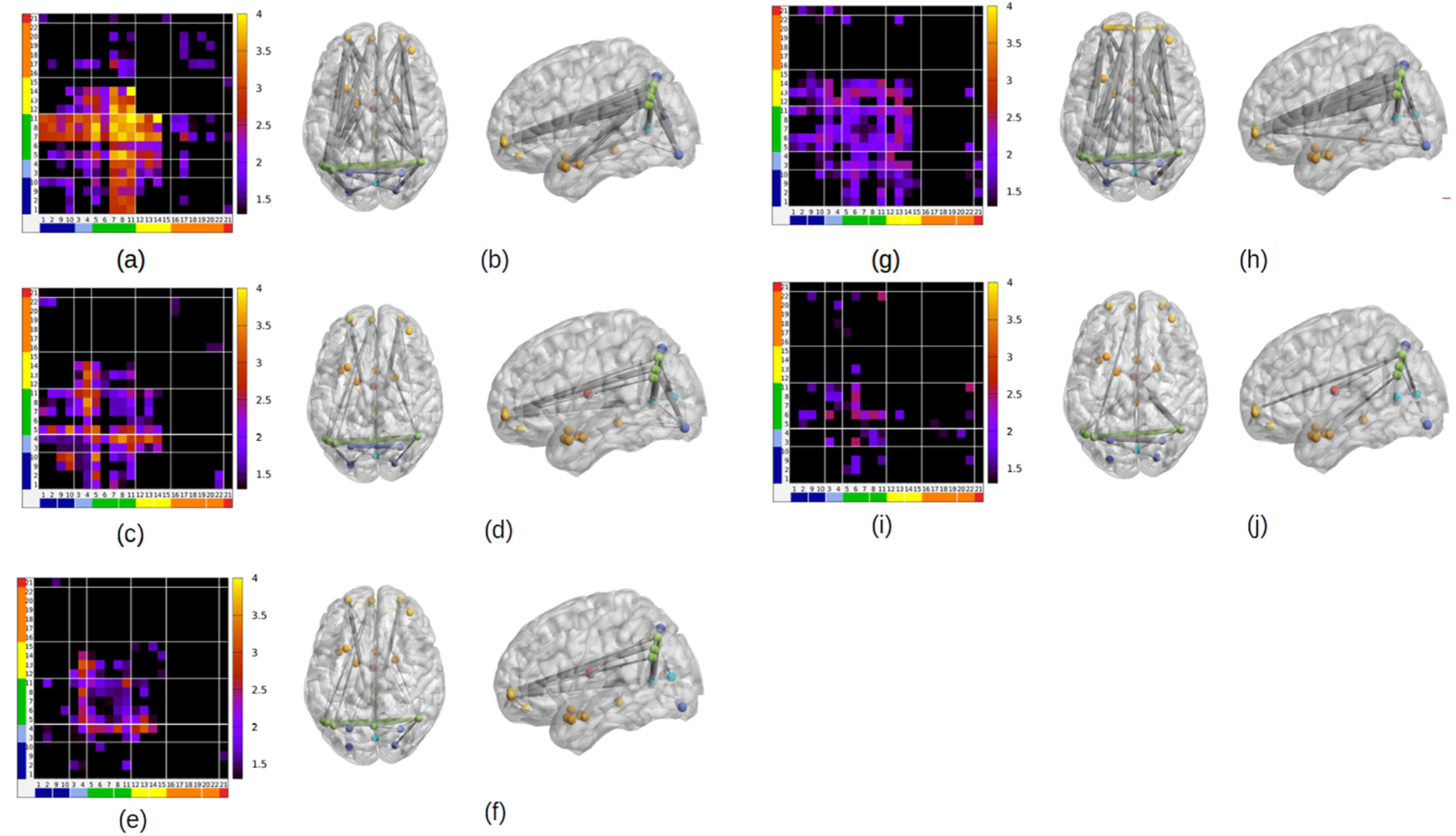
**(a-b)** Connectivity increase in blocks 1–20 for spatial strategy users. The matrix in panel (a) shows the log *p*-value of the increase between blocks 1 and 20 (Wilcoxon test). Panel (b) shows the links with a significant increase (*p* < .05 uncorrected) **(c-d)** Connectivity decrease between block 20 and block 21 for spatial strategy users. The matrix in panel (c) shows the p-value of the increase between block 20 and block 21 (Wilcoxon test). Panel (d) shows the links with a significant decrease (*p* < .05 uncorrected). **(e-f)** Connectivity increase between blocks 21 and 24 for spatial strategy users. The matrix in panel (e) shows the p-value of the increase between blocks 21 and 24 (Wilcoxon test). Panel (f) shows the links with a significant increase (*p* < .05 uncorrected) **(g-h)** Connectivity increase between block 1 and the block when subjects switched strategy (block “-1”) for color strategy users. The matrix in the panel (g) shows the p-value of the increase between blocks 1 and −1 (Wilcoxon test comparing). Panel (h) shows the links with a significant increase (*p* < .05 uncorrected) **(i-j)** Connectivity decrease between the block when subjects switched strategy (block “-1”) and the subsequent block (block “+1”) for color strategy users. The matrix in the panel (i) shows the *p*-value of the decrease between blocks −1 and +1 (Wilcoxon test). Panel (j) shows the links with a significant increase (*p* < .05 uncorrected).

**Figure 12.**
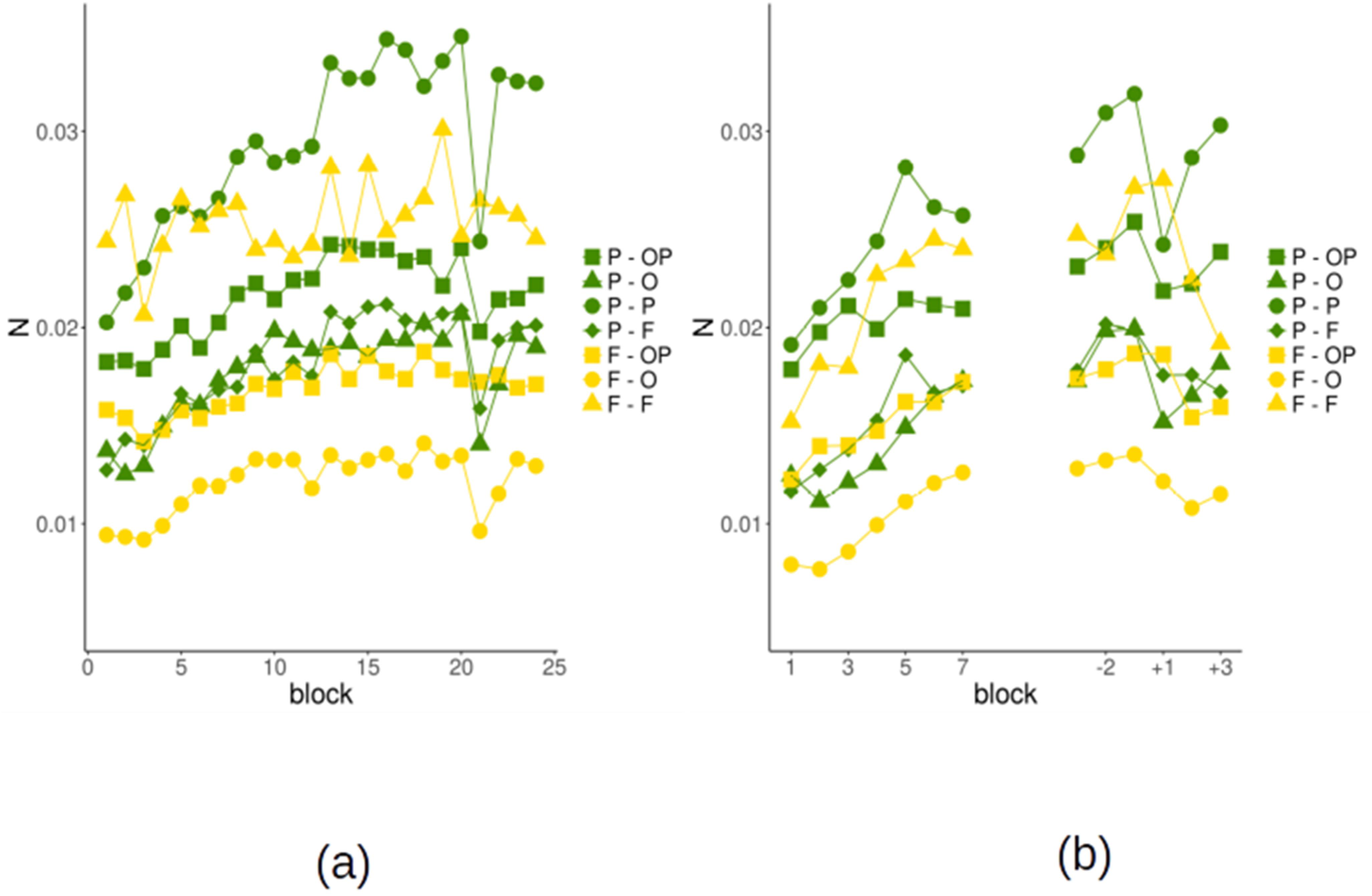
Strength of the links between modules as a function of the block for corner and color users. We show only links that have a significant effect (increase or change after the strategy shift) for either corner or color users (see Table 3). To facilitate the inspection of results, we show in green all links involving the parietal module, which have a similar behavior, and in yellow those involving the frontal module. **(a)** Strength of the links between modules as a function of the block for spatial strategy users. **(b)** Strength of the links between modules as a function of the block for color users. We show the 6 blocks around the switch after realigning the time series of each subject to the individual switch blocks (−1 is the block in which the subject spontaneously switches strategy, +1 the subsequent block).

We performed similar analyses for color users (Fig. 11d-e, 12b) by realigning the time series to the switch block. Overall the pattern is similar to the one found in spatial strategy users. Notably, however, color users showed an increase in fronto-frontal connectivity that was not present in spatial strategy users: Connectivity increased within medial prefrontal cortex (region 13) and between medial prefrontal and the two anterior lateral frontal regions bilaterally (region 12 and 14). Using the module-wise connectivity, we found a significant increase in P-P, P-OP, P-F, O-F, OP-F and F-F links between block 1 and the transition block (*p* < .05 FDR corrected, Wilcoxon test). Notably, the increase was significant even before the color-corner correlation was introduced (*p* < .05 FDR corrected, Wilcoxon test). Thus, color users seem to be characterized by a greater integration between regions of the F module, and between the F module and the occipito-parietal modules. Finally, in the passage to color strategy (from the transition block to the subsequent block) we observe a weak decrease of connectivity within the P module (*p* = .007 uncorrected), and between the latter and the F, OP and O modules (*p* =.06, *p* = .07, *p* =.06 uncorrected).

## Discussion

Humans can improve their performance in any task by gradually optimizing the implementation of a known strategy, or by devising and then adopting novel, more efficient strategies (Badre et al., 2010; Collins & Frank, 2013; Donoso et al., 2014; Heathcote et al., 2000; Roeder & Ashby, 2016; Schuck et al., 2015). Previous research has shown that practicing a task induces changes not only in the activation level of specific brain regions, but also in the long-range organisation of the relevant brain networks (Bassett & Mattar, 2017; Bassett et al., 2015; Chein & Schneider, 2005; Cole et al., 2013; Patel et al., 2013). However, the network dynamics governing strategy optimization versus the discovery of a new strategy are still unknown. By applying an analysis approach integrating the Coherence Density Peak Clustering (CDPC) (Allegra et al., 2017) with Multi-Voxel Pattern analysis, standard GLM and behavioral analysis, we could identify the brain regions involved in the task learning and performance. We showed that strategy optimization was specifically associated to an increase in local coherence and network integration centered on the precuneus and the angular gyrus bilaterally, corresponding to the parietal portions of the default network. By contrast, new strategy discovery was anticipated by a higher coherence and network integration in the medial and rostrolateral prefrontal cortex.

### Relevant brain regions: a fronto-parieto-occipital network

We identified 22 brain regions displaying high average local coherence over the ~1h of task performance in all subjects. A CDPC-based connectivity analysis showed that these 22 regions segregated in two main networks. A larger network involved frontal, parietal and occipital regions. This network could be further subdivided into smaller modules with a higher degree of internal coherence (see below). Another smaller network mainly involved temporal regions, and it did not exhibit a finer subnetwork structure (Fig. 10). High local coherence alone does not entail the involvement in a task: CDPC is an unsupervised approach, therefore it detects the presence of spatial and temporal coherence even when this is not task-related. However, converging evidence shows that at least the regions in the larger fronto-parieto-occipital network are indeed task-related. First, critical features of the stimuli, such as the spatial position, are represented in activation patterns in the regions of the network, as shown by a region-based MVPA analysis (Fig. 5). This finding is further supported by the overlap with regions reported previously using searchlight-based MVPA (Schuck et al., 2015). Second, all regions of the network are either activated or deactivated during task execution, as compared to a low-level baseline. Third, the local coherence and connectivity are strongly modulated by strategy optimization, strategy discovery, and strategy shift. By contrast, the other smaller network, mainly involving the temporal lobe, showed only limited converging evidence of being task-related. This leaves its functional role in this task open to interpretation. We focus our following discussion on the main occipito-parieto-frontal network.

### Incremental task optimization and instructed strategy change

The first aim of the present paper was to elucidate how coherence and connectivity vary when an established strategy is improved and when a shift to a new strategy occurs. Twenty-five out of thirty-six subjects continued to use the instructed spatial strategy and steadily improved their performance for ~50 minutes until they were explicitly told to switch to the color strategy for the last ~10 minutes. The optimization of the spatial strategy was associated with a progressive increase of local coherence in the precuneus, the lateral parietal lobe and in the medial occipital lobe. The increase of coherence in these regions was correlated with the reduction of reaction times during the optimization of the spatial strategy. When subjects were finally instructed to apply the color strategy, local coherence sharply dropped, as it would be expected if those regions were indeed involved in spatial strategy optimization. In the course of the instructed application of the new color strategy, coherence showed a fast recovery returning to the pre-switch level after only one block (2.5 minutes) in the anterior calcarine sulcus, in the precuneus and the angular gyrus bilaterally, thus suggesting that these regions were also involved when applying the new color strategy. While increasing their local coherence, these parietal and occipital regions showed either a decreasing or constant average activation. Furthermore, all occipital and parietal regions (with the exception of the anterior calcarine sulcus, see below) encoded the relevant spatial information of the stimulus. Altogether, these findings suggest that the progressive optimization of task processing shapes the activity of neural populations to become more coherent, specifically in regions involved in task processing. To the best of our knowledge, such relation between local coherence, learning, and task performance has not been reported yet (for a related measure mainly applied in resting state analysis see Jiang & Zuo, 2016).

Our findings indicate that a modulation of local coherence correlates with improved task performance. At present, however, we can only speculate as for the neural basis of this phenomenon. A modulation of local coherence is possibly achieved by a competitive mechanism enhancing task-relevant signals while reducing unrelated signals (Aston-Jones & Cohen, 2005; Eldar, Cohen, & Niv, 2013; Schmitz & Duncan, 2018). Furthermore, an increase in local coherence may indicate a progressive noise reduction, in line with recent neurophysiological findings. Works studying neuronal variability across multiple trials as a measure of the internal noise of a neural system have shown that the variability of task-relevant neurons decreases when stimuli are attended or perceived (Broday-Dvir, Grossman, Furman-Haran, & Malach, 2018; Churchland et al., 2010; Hussar & Pasternak, 2010; Mitchell, Sundberg, & Reynolds, 2007; Nougaret & Genovesio, 2018; Schurger, Sarigiannidis, Naccache, Sitt, & Dehaene, 2015). The reduction in neuronal variability has been associated to individual differences in perceptual ability, and to training in a working memory task (Arazi, Censor, & Dinstein, 2017; Qi & Constantinidis, 2012).

The findings on the four occipital regions we identified deserve a further comment. As mentioned above, the two regions in the anterior calcarine sulcus (3–4, Fig. 4) did not encode task-relevant information, while they showed learning effects similar to parietal regions. By contrast, the two regions in the posterior calcarine sulcus and lateral occipital cortex did not show learning effects but they encoded spatial information. Notably, posterior calcarine was activated compared to baseline while anterior calcarine was deactivated. We interpret this pattern as the effect of an attentional negative modulation on the portion of the calcarine sulcus processing the peripheral visual field, which is not useful for the task at hand. Thus, negative attentional modulation produces both a deactivation (Broday-Dvir et al., 2018) and a progressive increase in local coherence. By contrast, central visual field regions are activated, but their coherence already high at the beginning does not increase more with learning.

The analysis of the connectivity between regions and its dynamics provided both a confirmation of the findings on local coherence and further insights. The connectivity matrix over the whole experiment allowed to aggregate regions in few modules (Blondel et al., 2008; Sporns & Betzel, 2016) with stronger connectivity between regions within the module. Four modules were identified in the fronto-parieto-occipital network. Modules did not strictly follow mere anatomical proximity or functional subdivisions at rest (Power et al., 2011; Yeo et al., 2011). All frontal regions were in one module, but parietal regions and occipital regions were split. The module including regions in the angular gyrus and precuneus was largely composed by voxels belonging to the default network, similarly, the module including two occipital regions is entirely composed by voxels from the visual network. By contrast, the other two modules were mixed, one (the occipito-parietal) including both regions from the visual and the dorsal attention network, the other (frontal) including both voxels from the default network (the medial part of the network) and from the fronto-parietal control network (the lateral part). These departures from network organisation at rest underlie that the engagement in a task and practice shape the network organization of the brain (Power et al., 2011; Spadone et al., 2015; Yeo et al., 2011).

The connectivity dynamics followed a pattern similar to that of local coherence, particularly in the parietal module (“P” in Fig. 10). During the task optimization phase, the regions belonging to the parietal module greatly increased both the intra-module connectivity and the connectivity with all the other modules. Also, the connectivity showed a sharp drop upon strategy change. By contrast, other modules did not generally show a systematic increase of connectivity.

Overall, our findings suggest that the increase in regional and long-range connectivity is a driver of learning. An increased connectivity favors transfer of information and integration between brain regions, with possibly different functional specializations, but all involved in processing a task (Deco & Kringelbach, 2016; Shine & Poldrack, 2017). Compatible effects have been reported when comparing brain connectivity during rest to visuospatial attention (Al-Aidroos, Said, & Turk-Browne, 2012; Spadone et al., 2015), working memory (Cohen & D’Esposito, 2016; Shine et al., 2016) or flexible rule application (Vatansever, Menon, & Stamatakis, 2017). By contrast, the evidence on the network dynamics associated with learning is still limited, also because the analysis tools available until recently had low sensitivity in detecting network changes (Bassett & Mattar, 2017; Bassett et al., 2015). One important study by Bassett and collaborators reported that visuomotor learning was associated with increased functional segregation (i.e. decreased temporal coherence Bassett et al., 2015), a finding seemingly in contrast with ours. We argue that this diversity is due to the different processing requirements of the tasks explored in the two studies. The task we considered required the application of simple visuomotor rules implying the trial-by-trial integration of visual features to produce the appropriate motor response. Even when the task was highly practiced, subjects needed to rely on visual information to produce the correct response. In the study by Bassett and collaborators, by contrast, subjects were required to produce motor sequences, that once recognized and learned, could be generated from memory without further relying on visual information. This may explain why learning produced a segregation of the visual and motor network (see also Cohen & D’Esposito, 2016). Furthermore, in the study by Bassett and collaborators training was much longer (many hours over 42 days compared to 50 minutes in our case), possibly explaining the release from “control hubs”.

Notwithstanding the lack of network segregation during learning, the observed connectivity increase was far from homogeneous. The parietal module, a part of the default network, had a central role, being the only one increasing connectivity with all other modules. This finding highlights an active role of the default network during task processing, in contrast with the idea that the default network is shut down when a subject is engaged in a task (Crittenden, Mitchell, & Duncan, 2015; Margulies et al., 2016; Spreng, 2012; Vatansever, Menon, Manktelow, Sahakian, & Stamatakis, 2015). Thanks to its widespread connectivity, as emerged also in the present experiment, the default network could both receive sensory information and affect all task-relevant regions, to optimize stimulus processing and decision (Bar, 2007; Margulies et al., 2016). This possibility is consistent with recent works emphasizing the role of the default network also in optimizing and automatizing information processing (Dohmatob, Dumas, & Bzdok, 2018; Vatansever et al., 2017). Interestingly, the fact that the intra- and extra-module connectivity of the parietal module rebounds one block after switch suggests that optimization is an abstract process, recruited for strategies as diverse as those based on spatial information and color information. By contrast, during switching, when the system is reorganizing to cope with the new strategy the optimization is transiently paused.

### Spontaneous alternative strategy discovery and change

The second aim of the present paper was to understand how the spontaneous generation of a new strategy is related to coherence and connectivity dynamics. Eleven subjects discovered the uninstructed color strategy and applied it at a variable moment during task performance. These subjects showed an overall coherence and connectivity dynamics similar to the one described for corner strategy users, but with revealing differences. Similar to spatial strategy users, color strategy users showed an increased local coherence in the occipital and parietal regions before the spontaneous strategy change. Importantly, however, only color users showed an increase of local coherence in the anterior prefrontal regions. This specificity in local coherence was mirrored also in connectivity. While the intra-module connectivity of the prefrontal module remained constant in spatial strategy users, in color users the connectivity increased (Fig. 11). Notably, the tendency to increase started immediately, even before the color and the response were associated (first 4 blocks). Moreover, as spatial strategy users, color users showed a switch-related drop in local coherence and connectivity in the parietal module.

Overall, these findings suggest that connectivity patterns in medial prefrontal and rostrolateral prefrontal cortex reflect processes involved in the spontaneous discovery of novel strategies. Notably, connectivity among different frontal regions differed between participants long before their behavior began to change, foreshadowing who will discover a novel strategy and who will not. Rostrolateral prefrontal cortex has been proposed to be responsible for the evaluation of potential alternative strategies (Badre & Nee, 2018; Domenech & Koechlin, 2015; Donoso et al., 2014), while our own work has suggested that medial prefrontal cortex is involved in the internal simulation of an alternative strategy (Schuck et al., 2015). Moreover, the frontal regions also involve parts of orbitofrontal cortex that has been linked to the representation of task states, i.e. the information based upon which choices are currently selected (Badre & Nee, 2018; Schuck, Cai, Wilson, & Niv, 2016; Wilson, Takahashi, Schoenbaum, & Niv, 2014). It is thus possible that the connectivity increases reflect cross-talk between the above-named computations that are involved in finding and implementing a novel strategy. While this is also consistent with proposals relating the default network to background exploration (Bar, 2007; Constantinescu, O’Reilly, & Behrens, 2016; Crittenden et al., 2015; Dohmatob et al., 2018; Margulies et al., 2016; Vatansever et al., 2015), our findings additionally suggest a functional differentiation within the default network (Karahanoğlu & Van De Ville, 2015). The observation that the dynamics in the two subjects groups diverged from the beginning of the experiment suggests that the equilibrium between these two poles might be a relatively stable individual feature (Beaty et al., 2018; Melnick, Harrison, Park, Bennetto, & Tadin, 2013).

## Conclusions

In this study, we explored how brain networks behaved while human subjects optimized their strategy for solving a task or created a new one. The observed network dynamics suggested a pivotal role of default-mode network regions, but with a clear functional differentiation within the network. While the posterior part of the default-mode network increased connectivity and local coherence when subjects optimized their current strategy, the anterior part together with the rostrolateral prefrontal portion of the network was only involved in subjects who changed strategy. We speculate that the partially different behavior of the default-mode network in different persons might be a stable individual feature.

## Methods

### Task

Behavioral and imaging data of the main experiment were recorded while participants performed a simple perceptual decision-making task (Spontaneous Strategy Switch Task, Schuck et al., 2015). Participants were instructed to respond manually to the position of a patch of colored dots within a square reference frame. They were asked to select one of two responses depending on which corner of the reference frame the colored squares were closest to. Participants held a button box in each hand and could press either left or right. Two opposite corners (along with the diagonal) were mapped to the same response. The main task during scanning included twelve runs with 168 trials each. In Runs 1 and 2 (Random Runs), the stimulus color was unrelated to the position of the stimulus and the response. In Runs 3–10 (Correlated Runs) the color had a fixed relation to the response (e.g., all upper-left and lower-right stimuli were green, the remaining ones were red). Participants were not informed about this contingency but could learn and apply it spontaneously. By the end of Run 10, all participants were informed about the existence of a fixed association between color and corner (without specifying the relation) and instructed to use the color from then on (Instructed Runs). Each of the twelve runs of the main experiment lasted about 5 min and was followed by a short break. The experimenter monitored the performance of the participants. Written and oral feedback was given between runs if the error rate exceeded 20%. The response-stimulus interval was 400 ms. To measure the learning and use of color information, different trial conditions were used (for details, see Schuck et al., 2015). In the standard condition (80 out of 168 trials/run), the patch of dots was presented for 400 ms and was closest to one of the four corners of the reference frame; in the ambiguous trials (32 out of 168) the stimulus was centered within the reference frame and was presented for 400 ms; in the NoGo trials (32 out of 168) the colored squares were displayed for 2,000 ms without a reference frame in some trials and the task afterward continued with the next trial, with participants having to hold back any key press on the current trial; in the LateGo trials (16 out of 168), the frame was displayed after the initial 2,000 ms, and the participants had to react in a regular fashion; finally, in eight trials of each run the screen remained black for 3,000 ms (baseline condition). Due to the duration of the hemodynamic response function, the fast design of the experiment resulted in event-related BOLD signals, which also contained a signal proportion that reflected brain activation caused by previous and following events.

Before entering the scanner, participants were instructed and trained in the task. The instructions described all conditions (except ambiguous trials). Participants were only told to press any key of their choosing in case they were uncertain about the stimulus location. The color of the stimuli was mentioned only in an unspecified manner (“A stimulus can be either red or green.”). The response mapping was shown in all color combinations (a stimulus in each of the four corners was shown in both red and green during the instruction). In the training phase, participants were slowly accustomed to the short display durations (the display duration was successively shortened until it reached 400 ms). Feedback was given for all wrong and premature responses and time-outs (2,500 ms threshold). The color of the stimuli had no systematic relation to stimulus position during training. The training lasted at least 50 trials and ended when the participant made less than 20% errors in 24 consecutive trials. If the participant exceeded 168 trials without reaching the criterion, the training was restarted. Participants were further instructed that upon entering the scanner, no more feedback would be provided. After completion of the main experiment, participants completed a questionnaire with the following questions: (1) “In the experiment, which you have just completed, each corner had one associated color. Did you notice this while you were performing the task?” [yes/no]. (1b) “If yes, when did you notice this (after what percentage of the experiment)” [participants had to mark their answer on a scale from 0% to 100%]. (1c) “Did you use this color-corner relation to perform the task, i.e. to choose which button to press?” [yes/no]. (2) “Please indicate now which color the stimulus had for each of the four corners. If you did not notice this relation during the experiment or you are uncertain, you can guess.”

### Clustering frequency maps

Coherence Density Peak Clustering (CDPC, Allegra et al., 2017) aims at finding groups of voxels (*clusters*) whose BOLD signal is coherent in a given time window, usually short (e.g. 20 seconds). Contrary to other methods, such as community partition on a connectivity matrix, CPDC does not simply split all voxels into different clusters. In fact, voxels can be poorly coherent with other voxels (and hence not clearly part of any well-defined cluster), or coherent with other voxels only as a consequence of correlated fluctuations in the noise (a spurious cluster). CDPC first discards all voxels displaying poor or potentially artifactual coherence and then assigns only the remaining voxels (usually a small fraction of the total) to clusters.

The method starts by defining a *distance d_ij_* that captures the coherence between the BOLD signals of voxels *i* and *j*. The distance is given by the Euclidean distance between the BOLD time-series of the two voxels

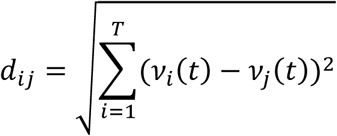

where, however, the raw time-series *v_i_(t)*, *v_j_(t)* have been suitably pre-processed, undergoing demeaning and amplitude-normalization. Note that the lowest frequency affecting the distances is 1/T (where T is the time window length), which for T = 22 s is .045 Hz.

Two voxels are regarded as coherent if the distance between the respective BOLD signals is low, as defined by a threshold, *d_ij_* < *d_c_*. Coherence between voxels can occur even if only noise is present. However, when only noise is present, high coherence tends to be observed between isolated voxels, while the presence of several coherent voxels within a close spatial neighborhood is unlikely (Allegra et al., 2017). More formally: for each voxel *i*, we define its neighbors as the voxels falling within a sphere of 6 mm radius centered on *i*, corresponding to about 27 voxels. We denote with *n_i_* the number of neighbors that are coherent with *i*. In our previous work, we showed that *n_i_* was generally lower than a threshold *n_0_* = 4 when only noise is present. All voxels with *n_i_* ≥ *n_0_* are thus considered in the clustering procedure, while voxels with *n_i_* < *n_0_* are discarded. This filtering procedure was shown to minimize the rate of detection of spurious clusters (Allegra et al., 2017). This criterion of cluster membership relies entirely on a measure of coherence that is strictly spatially local (27 neighboring voxels).

We run this procedure on sliding windows of 11 scans (22 s). This is the same length for which CDPC was validated (Allegra et al., 2017), and it is considered as the minimal window length for which transient connectivity clusters can be reliably identified (Hutchison et al., 2013). We use overlapping windows, progressively shifting the center of the window by 1 scan. The procedure is applied twice for each subject, the first time including only grey matter voxels, and the second time only white matter voxels.

For each subject, and for each voxel *i*, the clustering yields a binary value for every time window *t*, tracking whether the voxels was or not in a cluster. We devised an index, Φ, measuring how often a voxel *i* is part of a cluster in an interval comprising *N_t_* time windows.

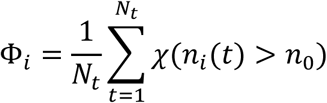

where *t* is a time window label, *N_t_* the number of time windows, *χ* is a step function (*χ*(*n_i_*(*t*) > *n_0_*) = 1 if *χ*(*n_i_*(*t*) > *n_0_*), *χ*(*n_i_*(*t*) > *n_0_*) = 0 otherwise). Intuitively, Φ_*i*_ is an aggregate measure of the coherence of the local activity of a voxel with its surroundings. We call Φ_*i*_ *clustering frequency* map, since, as we discuss below, if voxel satisfies the condition *n_i_*(*t*) > *n*_0_ it is automatically included in one of the clusters (see (Allegra et al., 2017)).

We compute a clustering frequency map for each *block*, i.e. half of a run (~150 s, Schuck et al., 2015). Thus for every subject, we generated 24 maps. The information given by a Φ map is not equivalent to the one obtained by running CDPC on a single time window equal to the entire block. The latter choice would include and emphasize the contribution of low frequencies (.005 Hz < f < .05 Hz) in the computation of *d_ij_*. This would reduce the sensitivity of the procedure to higher frequencies (f > .05 Hz), which are likely those critical for capturing the task related signals. The Φ maps focus on transient coherence occurring over timescales shorter than the whole block (see also Sakoğlu et al., 2010).

### High-coherence regions

The Φ_*i*_ maps can be used to identify in a data-driven way voxels that are potentially relevant for a task, under the assumption that task relevant voxels would be more often part of a cluster than voxels that are not (Allegra et al., 2017). For each subject, we averaged Φ maps over all blocks, obtaining one map for each subject. We normalized the average individual maps to MNI space and performed a Gaussian smoothing (FWHM = 9 mm). Finally, we averaged individual maps to obtain a single group map 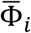 representing, for each voxel, the probability of being part of a cluster during task execution over all subjects. We assumed that coherence observed in the white matter could be safely ascribed to physiological and non-physiological noise. Therefore, we could use the distribution of 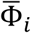 in white matter to identify a threshold above which Φ_*i*_ would be unlikely produced in the absence of a real (e.g. task-related) signal. We repeated the procedure described above, this time including in the analysis only white-matter voxels. We conservatively used as threshold the maximum value 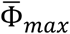 found in the white matter. Intuitively, the resulting thresholded map of the grey matter represents the voxels that are likely involved in the task.

To study the dynamics of Φ, we aggregated voxels above the 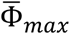 threshold in regions around every peak in the Φ map. The detailed procedure is the following:

a. We rank the voxels in order of decreasing 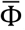;
b. We loop over all voxels above the threshold, starting from the voxel with the highest 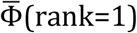. At iteration *n*, we check if the voxels contiguous to the voxel of rank n have been already assigned. If not, then all contiguous voxels must have lower values of 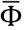: the voxel of rank *n* is hence a peak and starts a new region. If instead some of the contiguous voxels have already been assigned, it implies that they have higher values of 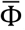, and we assign the voxel of rank n to the same region of these voxels. In case of ambiguity, we assign it to the same region of the contiguous voxel with highest 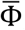. In this way, we could define regions tailored to the average spatial distribution of Φ.

### Connectivity: regional and long-range coherence

Φ maps identify voxels that are frequently coherent with their close spatial neighbors, and are thus assigned to clusters. Given that, voxels in the same high-Φ region can be considered, over the whole experiment, mutually coherent. However, Φ does not measure to which extent voxels within a high-Φ region, or in different high-Φ regions are mutually coherent. To answer this question, one needs to know not only whether two voxels are part of *any* cluster (as Φ does) but, more specifically, whether two voxels are part of the *same* cluster. To assign each voxel in each time window and in each subject to a specific cluster proceed as follows (Allegra et al., 2017).

We first define the *density ρ_i_* as the number of (non-isolated) voxels that are coherent with *i*, over the whole brain:

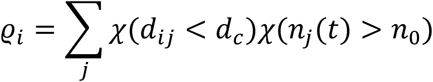

Notice that *ρ_i_* is usually higher than the number of coherent neighbors (measured by *n*_*i*_ for Φ) because typically a voxel is coherent with many voxels outside its local neighborhood. Cluster centers are identified as peaks in the density distribution. Following (Rodriguez & Laio, 2014), we compute 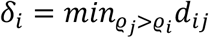, which is the minimum distance (in the space of BOLD signals) from a voxel higher density. Cluster centers stand out as isolated points with a large values of *δ_i_*. We rank the voxels according to their value of *δ_i_* and consider as putative cluster centers the first *k_max_* = 10 (Allegra et al., 2017). After the cluster centers have been chosen, all remaining voxels are assigned to a cluster following a recursive procedure. Each voxel is assigned to the same cluster of the most similar voxel having a higher density; if the latter voxel in not yet assigned, one looks for the voxel most similar to it having a higher density, and so forth until either an already assigned voxel or one of the cluster centers is reached. At the end of the procedure, we obtained a map, for each time window, assigning each voxel to a specific cluster.

Given two regions *a* and *b* we define a measure of their mutual coherence as

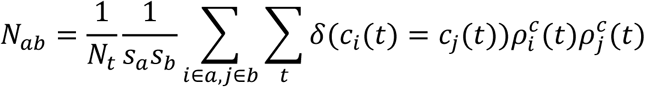

where *S_a(b)_* is the number of voxels in region a (resp. b). The term *δ*(*c_i_(t)* = *c_j_(t)*) is equal to one if voxel i and voxel j belong to the same cluster at time t. We weight this term by its density normalized to that of the cluster center, 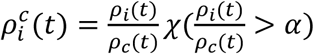 where *ρ_c_* (*t*) is the density of the cluster peak and *α* is a lower cutoff threshold (here, *α* = .3). In this way, we weight more pairs of voxels in the cluster *cores* (high density) than to the cluster *tails* (low density). The diagonal terms *N_aa_* measure the local coherence within the region *a*, while the off-diagonal terms *N_ab_* measure long-range coherence between different regions.

### Statistical tests

Given two group samples of Φ maps obtained in different conditions, we perform non-parametric statistical tests. When comparing the results for two disjoint groups of subjects we use a nonparametric statistic for independent samples - such as the Anderson-Darling statistic - to compare the distribution of Φ in the two samples; when comparing different conditions within a group of subjects, we can use a nonparametric statistic for dependent samples, such as the Wilcoxon signed-rank statistic. Indeed, Φ maps for different runs of the same subject can be correlated, so they cannot be assumed to be statistically independent.

Once the test statistic is chosen, we apply the test region-wise by averaging the Φ map over the regions, obtaining a region-wise map. In order to correct for multiple comparisons, we set in advance a threshold (usually, *α* = .05) and control the false discovery rate (FDR, Benjamini & Hochberg, 1995). In multiple tests with *N* null hypotheses, the FDR is defined as the expected ratio between the number of false rejections and the total number of rejections. In order to keep the FDR below a threshold *α*, one can follow a simple procedure. First, the p-values for each hypothesis are computed. Second, the p-values are sorted in increasing order, *p_1_* ≤…≤ *p_a_* ≤…≤ *p_N_* and one identifies *a*\*, the maximum *a* (if any) such that *p_a_* < *αa*/*N*. Finally, one rejects all hypotheses such that *p_a_* ≤ *p_a*_*.

We test for changes in Φ in selected regions. Since the regions are selected on the basis of the average Φ, hence on a criterion that is not independent of the data that are to be tested, in principle we might incur in a “double dipping” issue (Kriegeskorte, Simmons, Bellgowan, & Baker, 2009). In fact, we do not meet such a problem. A case of double-dipping would occur if, by selecting voxels with high average Φ across runs, we were inadvertently raising the chance of obtaining false positives. More formally, under the null hypothesis that the distribution of Φ_*i*_ across subjects is the same in the two different runs the statistic *W_i_* follows a well-defined distribution on which one computes *p*-values; a problem would occur if, by restricting attention to voxels with high average Φ, the resulting distribution *W_i_* were biased towards higher values of *W_i_*, inflating the likelihood of false positive detection. We can rule out this possibility by means of the following simulation, which shows that our selection criterion does not impact the null distribution of the chosen test statistics. Under the null hypothesis, the voxel-wise distributions Φ_*i*_ are the same in the two different runs. We can then generate two hypothetical samples Φ*_i,r1_* and Φ_*i,r2*_, corresponding to the samples of Φ_*i*_ obtained in two different runs *r*_1_, *r*_2_ across subjects, taking two Gaussian samples with mean given by the experimentally measured 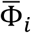 and variance given by 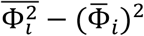. We then compute the distribution of *W_i_* over all voxels and over the 10% voxels with highest 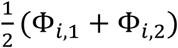 in the two runs, averaged over subjects. The latter subset corresponds to our selection criterions: restricting attention to voxels with a high average across runs and subjects. The distributions of *W_i_* are the same in the two cases (not shown). This proves that choosing voxels with high 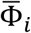 (averaged over subjects and runs) does not artificially inflate the detection of false positives, under the null hypothesis that the distribution of Φ is the same in different runs.

### Scanning and preprocessing

Acquisition of magnetic resonance images was conducted at the Berlin Center for Advanced Neuroimaging, Charité Berlin. We used a 3 T Siemens MagnetomTrio (Siemens) research-dedicated MRI scanner to acquire all data. T1-weighted structural images were acquired with an MP-RAGE pulse sequence with a resolution of 1 mm3. A T2*-weighted echo-planar imaging (EPI) pulse sequence was used for functional imaging (3 × 3 × 3 mm voxels, slice thickness = 3 mm, TR = 2,000 ms, TE = 30 ms, FOV = 192 mm, flip angle = 78°, 33 axial slices, descending acquisition). EPI slices were aligned to the anterior-posterior commissure axis. Field maps for distortion correction were acquired also using an EPI sequence. To allow for T1 equilibration effects, the experiment was started 6 s after the acquisition of the first volume of each run. Image pre-processing was performed using SPM12 (Wellcome Trust Centre for Neuroimaging, London, United Kingdom) running under Matlab 7.4 (R2007a) (Mathworks, Sherborn, MA, USA). The performed preprocessing steps were: a correction for magnetic inhomogeneities using field maps, slice timing correction, realignment to correct for motion, co-registration of anatomical images with functional images, and tissue segmentation based on the co-registered structural images to build a brain mask. Whenever required to allow for comparison of results across subjects, spatial normalization and/or smoothing was performed on the Φ maps (Allegra et al., 2017). The Φ maps were normalized to the standard MNI template and spatially smoothed with a Gaussian kernel of 9mm FWHM. For visualization purposes, we used the MRIcron software (www.mricron.com) and BrainNet Viewer (Xia et al., 2013; www.nitrc.org/projects/bnv/).

### GLM analysis

We performed a GLM analysis, with standard trials and resting trials as separate regressors, and motion parameters as nuisance variables. The response in standard trials was modeled as a response to stimulus presentation: onsets and durations corresponded to the onsets and durations of stimulus presentation. Resting trials had a duration of 3000 ms. We tested for significant activation or deactivations in the high-Φ regions, by averaging the contrast map over each region and performing a region-wise T-test over subjects.

### MVPA analysis

Representation of stimulus features (color and corner) was analyzed by a multivariate classification approach based on a support vector machine (SVM) with a linear kernel in combination with a searchlight approach (Haynes, 2015; Kriegeskorte, Goebel, & Bandettini, 2006; Norman, Polyn, Detre, & Haxby, 2006). For details, we refer to the previous work (Schuck et al., 2015). For color representation, the SVM was trained on parameter estimates (‘‘betas’’) from a general linear model of red and green NoGo trials in the last two runs (where all participants use the color strategy), and then tested on betas from Runs 1–10. This resulted in one accuracy map for each block and subject. For corner representation, the classifier was trained on betas of standard trials in the first two runs (where no participants use the color strategy) and then tested on betas from Runs 3–12.

## Acknowledgments

CR and SSA were supported by the PRIN grant 2010RP5RNM_001 from the Italian Ministry of University. The CDPC code used in this work can be downloaded from github (https://github.com/micheleallegra/CDPC).

